# FoTO1 is an epoxide isomerase in paclitaxel biosynthesis

**DOI:** 10.64898/2026.03.30.715439

**Authors:** Jie Bai, Jing Li, Yuanyuan Zhang, Hong Chang, Ningwei Zhang, Yuwan Liu, Jian Cheng, Xiaonan Liu, Huifeng Jiang

**Affiliations:** Tianjin Institute of Industrial Biotechnology, Chinese Academy of Sciences, Tianjin 300308, China; University of Chinese Academy of Sciences, Beijing 100049, China; State Key Laboratory of Elemento-Organic Chemistry, College of Chemistry and Frontiers Science Center for New Organic Matter, College of Chemistry, Nankai University, Tianjin 300350, China; Cooperative Innovation Center of Industrial Fermentation (Ministry of Education & Hubei Province), Key Laboratory of Fermentation Engineering (Ministry of Education), Hubei Key Laboratory of Industrial Microbiology, National “111” Center for Cellular Regulation and Molecular Pharmaceutics, Hubei University of Technology, Wuhan 430068, China; State Key Laboratory of Quality Research in Chinese Medicine, Institute of Chinese Medical Sciences, University of Macau, Macao SAR 999078, China; College of Biotechnology, Tianjin University of Science and Technology, Tianjin 300457, China

**Keywords:** FoTO1, Epoxide isomerase, Paclitaxel biosynthesis, QM/MM

## Abstract

Paclitaxel biosynthesis is limited by the instability of taxadiene-4(5)-epoxide, which readily diverts to the non-productive byproduct 5(12)-oxa-3(11)-cyclotaxane (OCT) instead of rearranging to taxadiene-5α-ol. Although FoTO1 suppresses OCT accumulation, its molecular function has been unclear. Here we identify FoTO1 as a dedicated epoxide isomerase that directs productive rearrangement. Biochemical characterization, site-directed mutagenesis, and QM/MM calculations reveal a pre-organized D68-D149 dyad that electrostatically activates epoxide ring opening and stereospecific rearrangement. Modular dissection of the C-terminal extension further reveals a functional partition between catalytic integrity and productive coupling with T5αOH, mediated by specific hydrophobic contacts that enforce precise geometric complementarity at the binary complex interface. These results demonstrate how electrostatic activation and enzyme association cooperate to control the fate of a highly reactive intermediate in paclitaxel biosynthesis.

## INTRODUCTION

Paclitaxel (Taxol) is a potent anticancer drug that has been widely applied in the treatment of diverse cancers^1,2^. The biosynthesis of paclitaxel in Taxus species relies on exceptional regio- and chemoselective control over complex diterpenoid intermediates^3-9^. A pivotal step in this pathway is the oxygenation of taxadiene catalyzed by the cytochrome P450 enzyme taxadiene-5α-hydroxylase (T5αOH)^4,10,11^. In fact, the direct oxidation product of taxadiene by T5αOH is taxadiene-4(5)-epoxide, rather than taxadiene-5α-ol (T5-ol) as previously proposed, since its active-site architecture lacks a suitable proton source to facilitate productive rearrangement of the epoxide^12^. Experimental characterization demonstrated that in acidic aqueous environments,taxadiene-4(5)-epoxide undergoes predominantly non-specific rearrangement to the non-natural byproduct 5(12)-oxa-3(11)-cyclotaxane (OCT), with only minor conversion to the biosynthetic target T5-ol^13^. This preferential rearrangement pathway poses a fundamental challenge to metabolic fidelity and suggests that Taxus species require an auxiliary catalytic component to govern the fate of this reactive intermediate.

The molecular identity of such an auxiliary factor remained elusive until the recent discovery of FoTO1 (facilitator of taxane oxidation) through transcriptomic correlation analysis in Taxus cell cultures^14^. FoTO1 was found to co-express with T5αOH and suppress OCT accumulation in heterologous expression systems. Microscale thermophoresis measurements further revealed that FoTO1 binds T5αOH with high affinity, suggesting a direct physical association^14^. However, the specific biochemical function of FoTO1 remained unclear. Whether FoTO1 possesses intrinsic catalytic activity to redirect epoxide rearrangement or serves primarily as a scaffolding element to facilitate T5αOH function represents a critical knowledge gap in understanding paclitaxel biosynthetic pathway.

In this work, we demonstrate that FoTO1 functions as a dedicated epoxide isomerase that governs the fate of the taxadiene-4(5)-epoxide intermediate through a dual mechanism. Combining *in vitro* reconstitution with chemically synthesized substrate, systematic mutagenesis, structural modeling, and QM/MM analysis, we identify a pre-organized D68-D149 dyad that creates an electrostatically optimized environment for epoxide ring-opening and stereospecific rearrangement. In addition, modular dissection of the FoTO1 C-terminal extension reveals a functional partition between catalytic integrity and productive coupling with T5αOH, mediated by specific hydrophobic contacts at the binary complex interface. Together, these results illustrate how catalytic activation and enzyme–enzyme organization cooperate to control a reactive intermediate in specialized plant metabolism and provide a framework for engineering metabolically robust biosynthetic pathways.

## RESULTS AND DISCUSSION

### FoTO1 Functions as a Specialized Epoxide Isomerase

FoTO1 belongs to the nuclear transport factor 2 (NTF2)-like protein superfamily, a structurally diverse group characterized by a conserved α/β-barrel fold that has been co-opted for a wide range of catalytic functions^15-17^. Within this superfamily, limonene-1,2-epoxide hydrolase (LEH) from *Rhodococcus erythropolis* represents a well-characterized epoxide-processing enzyme. LEH utilizes a catalytic network in which residues Y53 and N55 activate a nucleophilic water molecule, while D101 and D132 facilitate electrophilic activation and proton transfer^18^. Structural alignment of FoTO1 with LEH reveals conservation of overall active-site topology, with a closely preserved spatial arrangement of pocket-lining residues (Fig. 1a). However, detailed inspection of the FoTO1 catalytic center identifies key substitutions that alter the chemical environment. Although D149 in FoTO1 occupies a position analogous to D132 in LEH and is supported by a similar hydrogen-bonding network, the residues corresponding to the water-activating dyad (Y53/N55) and the electrophilic aspartate D101 are replaced by F64, D68, and Q118, respectively (Fig. 1a). Retention of the overall structural framework coupled with substitution of these catalytic residues suggests a divergence from the canonical hydrolytic mechanism.

**Figure 1.**
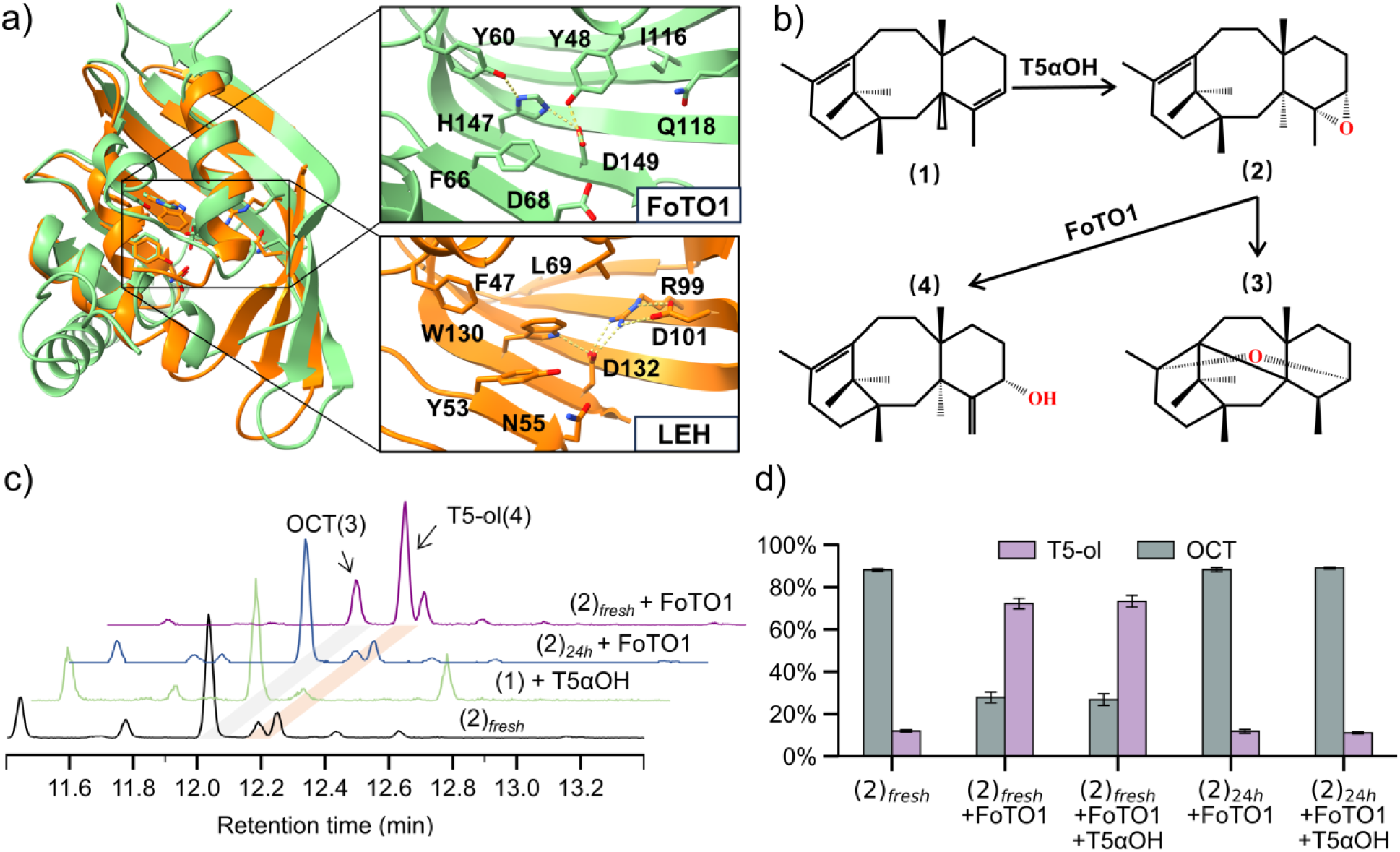
Functional characterization of FoTO1 as a specialized epoxide isomerase. (a) Structural overlay of LEH and FoTO1 α/β-barrel folds with labeled active-site residues. (b) Proposed processing pathways of taxadiene-4(5)-epoxide (2) to OCT (3) or taxadiene-5-ol (4). (c) Representative GC-MS chromatograms for the indicated reaction conditions. (d) Quantitative product distribution (OCT, grey; taxadiene-5-ol, purple) determined by GC-MS peak area. Data represent mean +/−SD (n = 3).

Our previous crystallographic and computational analyses established that T5αOH generates taxadiene-4(5)-epoxide (2) as its direct product upon oxygenation of taxadiene (1)^12^. However, the intrinsic chemical lability of this epoxide complicates its characterization and poses a challenge to metabolic fidelity. Although strong acid treatment can promote rearrangement of (2) to T5-ol (4), under non-enzymatic conditions that are more representative of physiological or analytical environments, including mildly acidic aqueous solutions or elevated temperatures, (2) predominantly undergoes non-specific rearrangement to the non-natural byproduct OCT (3). This instability diverts metabolic flux away from productive paclitaxel biosynthesis^13^. (Fig. 1b) Considering both its structural relationship to epoxide-processing enzymes and the intrinsic instability of (2), we hypothesized that FoTO1 acts as a dedicated isomerase that intercepts the T5αOH-generated epoxide. By providing a constrained catalytic environment, the enzyme was proposed to direct rearrangement toward (4) while limiting non-specific formation of (3) observed under non-enzymatic conditions.

To evaluate this hypothesis, chemically synthesized (2) was prepared using established epoxidation methods^13^. GC-MS analysis demonstrated that the spontaneous rearrangement profile of synthetic (2) in aqueous buffer (50 mM sodium acetate, pH 5.5, 30 °C) closely mirrors the product distribution observed during T5αOH-catalyzed oxygenation of taxadiene in the absence of downstream factors, with (3) as the predominant species. These data support the assignment of (2) as a physiologically relevant intermediate. The catalytic activity of FoTO1 was then examined *in vitro*. Co-incubation of purified FoTO1 with freshly prepared (2) in product distribution, resulted in a substantial increase in the proportion of (4) relative to buffer-only controls (Fig. 1c).

To determine whether FoTO1 acts on intact epoxide substrate or downstream rearrangement products, time-delay experiments were performed. Pre-incubation of (2) in buffer for 24 h prior to addition of FoTO1 led to increased formation of (3), consistent with spontaneous epoxide rearrangement (Fig. 1c). Under these conditions, FoTO1 did not restore the productive product distribution, indicating that the enzyme acts on the intact epoxide intermediate and does not reverse downstream rearrangement pathways. Supplementation with T5αOH did not alter the product profile relative to FoTO1 alone (Fig. 1d), indicating that the observed product redistribution arises from FoTO1 activity rather than residual P450 catalysis. Together, these results support a model in which FoTO1 competes with spontaneous epoxide rearrangement by intercepting (2) and directing its conversion toward (4).

### Catalytic Mechanisms of Epoxide Isomerization

To elucidate the molecular determinants of FoTO1-mediated isomerization, we performed systematic site-directed mutagenesis of active-site residues predicted by structural modeling to participate in the hydrogen-bond network, including Y48, Y60, D68, H83, H147, and D149 (Fig. 2a). Substitutions at Y60 and H83 yielded divergent outcomes. Variants Y60F, Y60A, and H83F abolished activity, whereas H83A retained near-wild-type function. Given that Y60 is deeply embedded within the hydrophobic core and forms stabilizing interactions with surrounding residues, the loss of activity in Y60F/A and H83F is more plausibly attributed to structural destabilization rather than direct catalytic involvement. The retained activity of H83A further indicates that the imidazole side chain at this position is not essential for catalysis and that the Y60-H83 hydrogen bond is unlikely to be mechanistically required. In contrast, substitutions at Y48, D68, H147, and D149 uniformly abolished activity across all tested variants (Y48F/A, D68N/A, H147F/A, D149N/A), identifying these residues as functionally indispensable. Structural analysis indicates that Y48, H147, and D149 form a pre-organized hydrogen-bond network that constrains the orientation of D149, with additional stabilization provided by Y60 through its interaction with H147.

**Figure 2.**
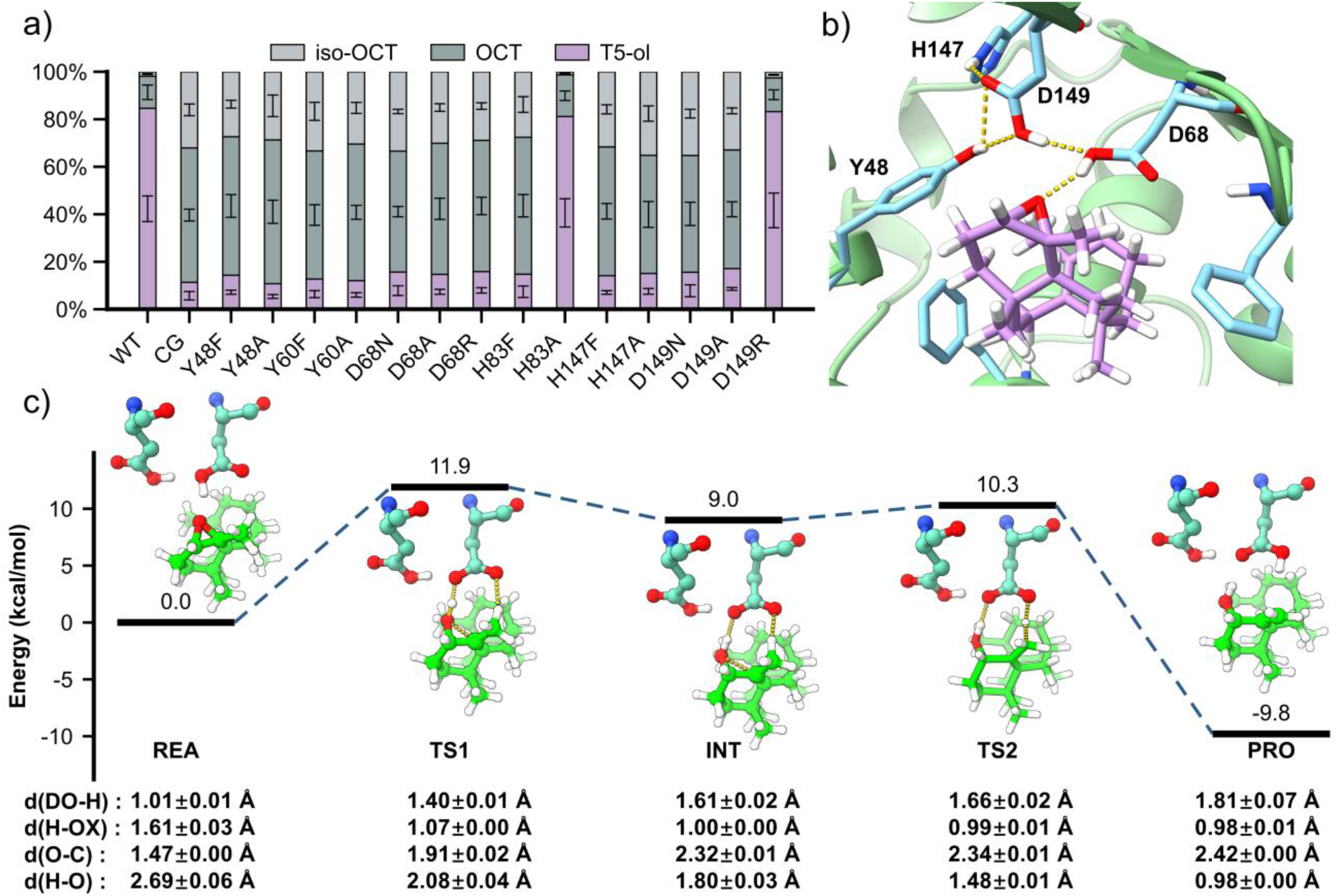
Mechanistic characterization of FoTO1-catalyzed epoxide isomerization. (a) Product distribution of FoTO1 active-site mutants in yeast co-expressing T5αOH. Ratios of iso-OCT, OCT and T5-ol were determined by GC-MS. Data represent mean +/−SD (n = 3). (b) Representative MD simulation snapshot of taxadiene-4(5)-epoxide (purple) binding. Catalytic residues (D68, D149) and the supporting hydrogen-bond network (Y60, H147, Y48) are labeled; yellow dashed lines denote hydrogen bonds. (c) Reaction potential energy surface (PES) calculated via QM/MM at the B3LYP-D4/def2-TZVP level. Relative energies are provided in kcal/mol. Insets show stationary point geometries with annotated interatomic distances (Angstroms).

A key mechanistic insight emerged from the D149R substitution. Introduction of a constitutive positive charge at this position restored catalytic activity to a level comparable to wild-type FoTO1, indicating that the critical feature at residue 149 is not hydrogen-bond donation per se, but the generation of a localized electropositive environment. In this context, the Y48-H147-D149 triad can be understood as serving a dual function: enforcing active-site pre-organization while modulating the electronic properties of the neighboring D68 residue. Accordingly, we propose that a D68-D149 catalytic dyad constitutes the functional core of the reaction. In this model, D68 serves as the primary proton donor for epoxide ring opening, while residue 149 modulates its proton-donating capability through electrostatic polarization rather than direct proton transfer (Scheme 1).

**Scheme 1.**
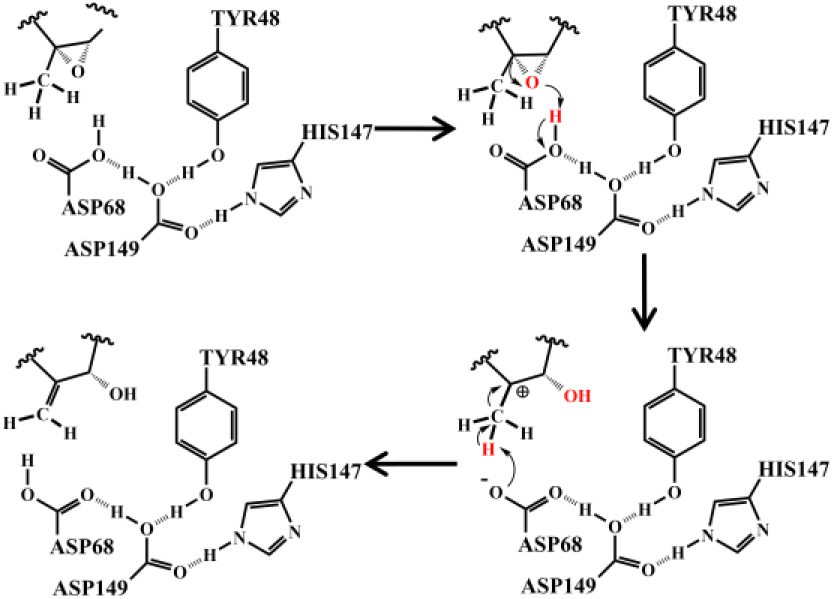
Proposed catalytic mechanism of FoTO1-mediated epoxide isomerization via an induced polarization strategy.

To evaluate the feasibility of this model, molecular docking and molecular dynamics (MD) simulations were performed on the FoTO1-substrate complex. The resulting structural ensemble indicates that D68 and D149 are embedded within an extensive hydrogen-bond network comprising Y60, H147, and Y48 (Fig. 2b). This framework constrains D68 in an axial orientation that positions its carboxyl proton for productive interaction with the epoxide oxygen. Hybrid QM/MM potential energy surface (PES) calculations were conducted to characterize the energetic landscape (Fig. 2c). The reaction is initiated by protonation of the epoxide oxygen by D68, which constitutes the highest-energy step along the computed pathway. This ring-opening transition state (TS1) exhibits an electronic barrier of 11.9 kcal/mol, leading to a carbocationic intermediate at 9.0 kcal/mol relative to the reactant state. Subsequent stereospecific deprotonation of the methyl group adjacent to the carbocationic center by D68 proceeds via a second transition state (TS2) at 10.3 kcal/mol, leading to double bond formation and yielding T5-ol, corresponding to a barrier of 1.3 kcal/mol relative to the intermediate. The disparity between these barriers suggests that once epoxide protonation occurs, the subsequent rearrangement proceeds with strong kinetic commitment toward T5-ol formation within the constrained active-site geometry. In contrast, an epoxide released into bulk aqueous solution lacks the geometric constraints imposed by the FoTO1 active site and is therefore susceptible to alternative acid-catalyzed rearrangements, consistent with the predominant formation of OCT observed under non-enzymatic conditions^12^. These findings suggest that the catalytic contribution of FoTO1 extends beyond rate enhancement to the imposition of geometric and electrostatic constraints that direct the rearrangement along a biosynthetically productive pathway.

To further characterize the electronic origin of this effect, we calculated the electronic electrostatic potential (EESP), defined as the electrostatic potential excluding nuclear contributions, at the position of the D68 carboxyl proton. For an isolated protonated carboxyl group, the calculated EESP value is −0.948 a.u. Within the native FoTO1 active site, where D149 forms an axial hydrogen bond with D68, the value shifts to −0.913 a.u. In the computational D149R model, the presence of the guanidinium group induces a further shift to −0.801 a.u. (Fig. S1). This systematic positive shift in EESP provides an electronic descriptor consistent with increased proton-donating propensity of D68 within the structured active-site environment. Importantly, the functional restoration observed in the D149R mutant corroborates this electrostatic polarization model, supporting the interpretation that residue 149 contributes primarily through modulation of the local electrostatic field rather than direct participation in proton transfer.

Collectively, mutagenesis, structural modeling, molecular dynamics simulations, QM/MM energy profiling, and electrostatic analyses support a unified mechanistic framework in which FoTO1 employs induced polarization to influence epoxide fate. By pre-organizing a D68-centered proton donor within a hydrogen-bond network and tuning its acidity through secondary-shell electrostatic effects, the enzyme creates geometric and electronic conditions that favor selective interception of the reactive epoxide intermediate. This integrated model provides a structural and energetic rationale for the preferential formation of T5-ol under enzymatic conditions.

### The Synergistic Catalysis Between FoTO1 and T5αOH

Having established the intrinsic isomerase activity of FoTO1, we next examined how its physical association with T5αOH contributes to metabolic flux control. Previous microscale thermophoresis studies demonstrated that FoTO1 binds T5αOH with high affinity^14^, suggesting a functional partnership beyond passive spatial proximity. To identify structural determinants responsible for the synergistic catalysis between FoTO1 and T5αOH, we performed a systematic modular dissection of the FoTO1 C-terminal extension, a domain unique to FoTO1 and absent in other NTF2-like family members.

Based on the distribution of localized polar and electrostatic clusters, the 41-residue C-terminal extension was partitioned into discrete functional modules, which were evaluated via a series of progressive truncations (Fig. 3a). These modules were evaluated using parallel *in vitro* biochemical assays and *in vivo* heterologous expression experiments to distinguish their respective contributions to intrinsic catalytic activity versus coupling efficiency. *In vitro* incubation assays employing synthetic taxadiene-4(5)-epoxide revealed that the proximal portion of the C-terminus (residues 21–41) is essential for maintaining catalytic competence. Truncation of the first 20 residues (Δ10, Δ20) did not impair conversion of the free epoxide to T5-ol, further removal of just four additional residues (Δ24) resulted in a marked attenuation of enzymatic activity, a deficit that persisted in the full C-terminal deletion (Δ41) (Fig. 3b). Structural analysis suggests that the C-terminal residues 21–41 forms a surface-exposed salt-bridge cluster that anchors the C-terminal extension to the globular NTF2-like core domain. Although not directly involved in catalysis, this cluster likely preserves the structural integrity of the catalytic chamber required for induced polarization.

**Figure 3.**
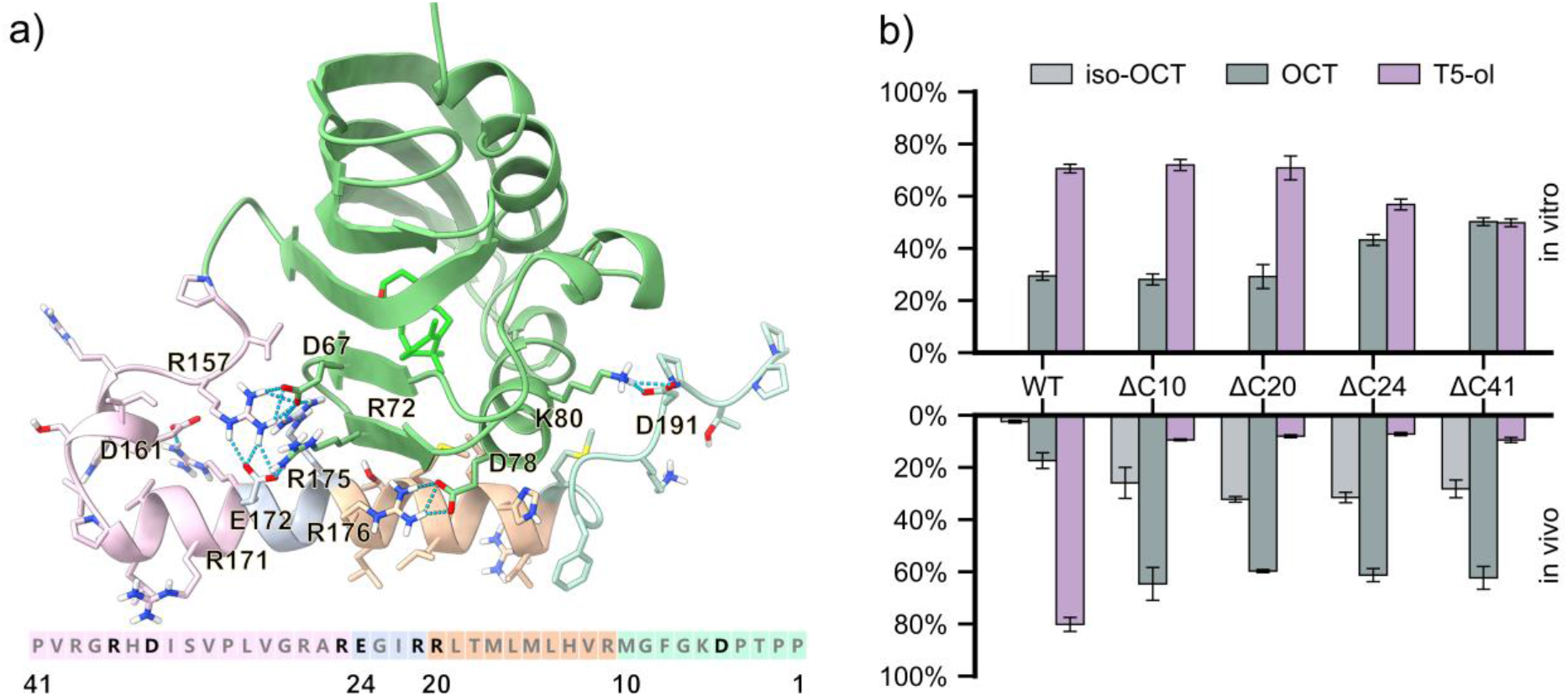
Structural modules of the FoTO1 C-terminus. (a) Partitioning of the FoTO1 C-terminal extension into four discrete segments. Salt-bridge residue pairs and sequences are indicated. (b) GC-MS product ratios for C-terminal truncations. Top: *In vitro* incubation with epoxide (2). Bottom: *In vivo* yeast fermentation with T5αOH.

In contrast, heterologous co-expression experiments in yeast revealed a functional decoupling between intrinsic catalytic activity and coupling efficiency. While the C-terminal residues 1–20 were dispensable for *in vitro* isomerase activity, their deletion in a cellular context caused metabolic flux to revert toward the spontaneous byproduct OCT, yielding product ratios resembling those observed in FoTO1-deficient strains (Fig. 3b). Together, these data reveal a functional partitioning within the C-terminal extension: the distal segment (residues 1–20) is required for efficient coupling *in vivo*, whereas the proximal framework (residues 21–41) is indispensable for stabilizing the enzyme’s intrinsic catalytic architecture.

### Mechanisms of the Coupling Between FoTO1 and T5αOH

To explore the structural basis of FoTO1-T5αOH coupling, we performed computational modeling of the binary complex using Protenix^19^. A total of 1000 independent predictions were generated, from which models exhibiting extensive interfacial contact between the FoTO1 C-terminal domain and T5αOH were selected for further analysis. Two distinct binding modes were observed, characterized by opposite orientations of FoTO1 relative to T5αOH. Despite these opposing orientations, both models converge on a common architectural feature: insertion of the FoTO1 C-terminal segments into a hydrophobic groove formed between the T5αOH N-terminal helix and the F/G helix pair, potentially shielding exposed hydrophobic surfaces and improving the solvent compatibility of the predicted complex (Fig. 4a, Fig. S5).

**Figure 4.**
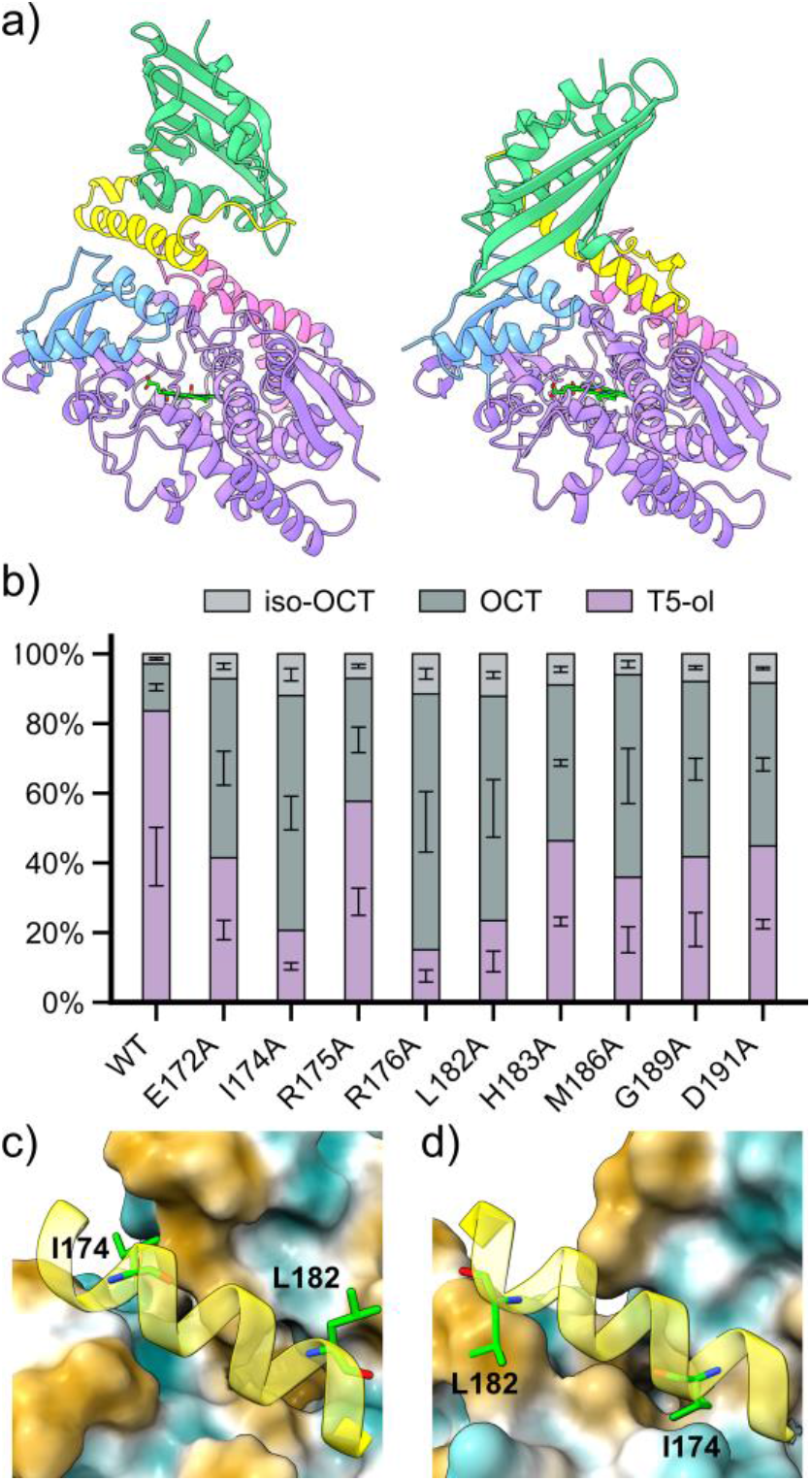
Structural modeling and mutational analysis of the FoTO1-T5αOH complex. (a) Predicted binding modes. FoTO1 core, green; C-terminus, yellow; T5αOH, purple; F/G helix, pink; N-terminal, blue. (b) Coupling efficiency of FoTO1 alanine scanning variants across residues 172–191, determined by GC-MS product distribution in yeast co-expressing T5αOH. Data represent mean ± SD (n = 3). (c, d) Structural details of the predicted binding interface. The segment bearing I174 and L182 of the FoTO1 C-terminal extension is shown in ribbon representation; T5αOH is displayed as a molecular surface colored by hydrophobicity (brown, hydrophobic; white, neutral). The interface-contributing residues I174 and L182 are shown as sticks and labeled.

To identify residues within the C-terminal extension that mediate productive interaction with T5αOH, we performed alanine scanning mutagenesis across C-terminal residues 1–24, guided by the predicted binding geometries. Nine positions were identified at which alanine substitution significantly reduced coupling efficiency relative to wild-type FoTO1 (Fig. 4b, FigS6). Interpretation of these variants was guided by cross-referencing the mutational outcomes with the predicted FoTO1-T5αOH interface. Although substitutions at E172, R175, R176, H183, and D191 reduced coupling efficiency, none of these residues were observed to form contacts with T5αOH in either predicted complex model. We therefore attribute their functional contribution to stabilization of the intramolecular architecture of the C-terminal extension relative to the NTF2-like core, rather than to direct intermolecular interaction. A similar interpretation applies to M186 and G189, whose alanine substitutions reduced coupling efficiency without evidence of direct contact with T5αOH, suggesting that these hydrophobic residues contribute primarily to intra-domain structural integrity rather than interface formation.

In contrast, substitutions at I174 and L182 reduced coupling efficiency under conditions where these residues are positioned at the predicted FoTO1-T5αOH binding interface. These findings are consistent with a model in which I174 and L182, residing on the hydrophobic face of the C-terminal extension, anchor productive binary complex formation by embedding within the hydrophobic groove of T5αOH, thereby enforcing the precise geometric alignment required for efficient intermediate capture (Fig. 4c, d). These findings demonstrate a requirement for precise geometric alignment between the T5αOH product exit channel and the FoTO1 substrate-binding pocket, mediated by specific hydrophobic contacts at the native C-terminal interface.

Collectively, these results define a two-tiered architectural specialization within the FoTO1 C-terminal extension. The proximal segment (C-terminal residues 21–41) preserves catalytic integrity through intra-domain structural contacts, whereas key residues within the distal segment, particularly I174 and L182, anchor productive association with T5αOH by embedding within its hydrophobic groove. This modular organization enables FoTO1 not only to catalyze stereospecific epoxide rearrangement, but also to confine the reactive intermediate within a geometrically defined enzymatic environment, thereby maintaining the coupling efficiency required for faithful early-stage paclitaxel biosynthesis.

## CONCLUSIONS

This study identifies FoTO1 as a dedicated epoxide isomerase that safeguards metabolic fidelity during early paclitaxel biosynthesis. Rather than merely accelerating a chemical rearrangement, FoTO1 functions as a control node that intercepts a chemically labile intermediate and enforces its productive conversion to T5-ol, thereby suppressing diversion into non-productive side products. These findings defined a dual-layer strategy for managing hyper-reactive intermediates in specialized metabolism: electrostatic steering to control reaction directionality, and geometric channeling to constrain intermediate diffusion. The modular organization of FoTO1, which separates catalytic tuning from interfacial docking, illustrates how enzymatic precision and supramolecular architecture can co-evolve to resolve metabolic bottlenecks imposed by unstable intermediates. More broadly, this work highlights that effective pathway engineering must consider not only catalytic efficiency, but also inter-enzyme geometry and electrostatic microenvironment as determinants of metabolic fidelity.

## MATERIALS AND EXPERIMENTAL METHODS

### Expression and purification of FoTO1 and T5αOH

The expression and purification methods of T5αOH were reported in previous literatures^20^. The nucleotide sequence of FoTO1 was optimized by E. coli codon optimization. The N-terminal of FoTO1 was ligated with MBP by a linker of six glycine residues and then constructed into pET28a vector (pET28a-MBP-FoTO1). The recombinant vector was transformed into *E. coli BL21 (DE3)* cells. A single colony was transferred to LB liquid medium containing 100 μg/mL kanamycin and cultured at 37 °C to an OD600 of approximately 0.6–0.8. FoTO1 expression was induced by addition of 0.3 mM IPTG and then cultured at 18 °C for 16 hours.

The cells were harvested by centrifugation at 6,000 g and resuspended in 30 mL lysis buffer (20 mM Tris-HCl (pH 8.0), 0.5 M NaCl, 10% glycerol, 30 mM imidazole). The cells suspensions were disrupted by a high-pressure homogenizer (JNBIO, China) and centrifuged at 15,000 rpm for 30 min at 4 °C to remove the cell debris. To bind the recombinant enzyme, which was expressed as a fusion protein containing a 6×His tag, the supernatant was filtered and loaded onto a Ni^2+^-chelating affinity chromatography column (GE Healthcare, USA) pre-equilibrated with lysis buffer. After the column was rinsed with 50 mL wash buffer (20 mM HEPES-KOH (pH 8.0), 0.5 M NaCl, 5% glycerol and 50 mM imidazole), the target proteins were eluted with 30 mL elution buffer (20 mM HEPES-NaOH (pH 8.0), 0.5 M NaCl, 5% glycerol and 250 mM imidazole). The eluted proteins were concentrated and dialyzed against dialysis buffer (10 mM HEPES-NaOH (pH 8.0), 50 mM NaCl) by ultrafiltration with an Amicon Ultra centrifugal filter device (Millipore, USA). The purity of proteins was evaluated by 4-20% SDS-PAGE. Protein concentrations were determined by BCA Protein Assay Kit (Pierce, Thermo Fisher Scientific, USA) using a microplate reader. Consequently, the proteins were stored at −80 °C after flash freezing in liquid nitrogen.

### *In vitro* reaction of FoTO1 and taxadiene-4(5)-epoxide

The substrate taxadiene-4(5)-epoxide was synthesized according to the method reported in previous literature^13^. 100 μL of the reaction system was carried out in 50 mM sodium acetate (pH 5.5) solution, including: 500 μM taxadiene-4(5)-epoxide and 1 mg/mL FoTO1. In addition, the reaction without FoTO1 and the reaction with 1 mg/mL T5αOH were used as controls. After reaction at 30 °C for 12h, the mixture was extracted with *n*-hexane and detected by GC-MS.

### *In vitro* validation of the C-terminal function of FoTO1

Using pET28a-MBP-FoTO1 as the template, the recombinant plasmids pET28a-MBP-FoTO1-t10, pET28a-MBP-FoTO1-t20, pET28a-MBP-FoTO1-t24, and pET28a-MBP-FoTO1-t41 were constructed, respectively. The sequence-verified plasmids were transformed into *E. coli BL21(DE3)* for protein expression and purification (following the procedures described above for FoTO1 protein expression and purification). The C-terminally truncated FoTO1 proteins of different lengths (FoTO1-t10, FoTO1-t20, FoTO1-t24, and FoTO1-t41) were individually incubated with taxadiene-4(5)-epoxide. The components of the 100 μL reaction system were the same as those used for the FoTO1 and taxadiene-4(5)-epoxide reaction described above. The wild-type FoTO1 protein and the reaction without protein addition were used as the positive and negative controls, respectively.

### Construction of *Saccharomyces cerevisiae* engineered strains and mutant experiments

The genes FoTO1, T5αOH, and TS were synthesized by GenScript (Nanjing, China) with codon optimization for S. cerevisiae. The Arabidopsis thaliana-derived P450 reductase ATR2 was amplified from the laboratory-preserved Y33-ATR2 vector. To optimize P450 expression, a recombinant Ycplac22 plasmid was constructed by linking the transmembrane-domain-truncated T5αOH with ATR2 under a bidirectional promoter via Gibson Assembly. Similarly, the TS gene (with a 24-bp N-terminal truncation) and FoTO1 were assembled into the Ycplac33 vector (Y33-MBP-tTS-FoTO1). All cloning procedures were performed using E. coli DMT. The resulting vectors were transformed into a high-efficiency GGPP-producing S. cerevisiae chassis (kindly provided by Prof. Xue-Li Zhang and Prof. Zhu-Bo Dai, TIB, CAS) using the lithium acetate method. Transformants were screened on appropriate synthetic dropout media at 30 °C.

### *In vivo* validation of the c-terminal function of FoTO1

Using Y33-MBP-tTS-FoTO1 as the template, the recombinant plasmids Y33-MBP-FoTO1-t10, Y33-MBP-FoTO1-t20, Y33-MBP-FoTO1-t24, and Y33-MBP-FoTO1-t41 were constructed, respectively. Each of the four sequence-verified plasmids was co-transformed with Y22-ATR2-t24T5αOH into the previously constructed *Saccharomyces cerevisiae* strain Sc-GGPP, yielding strains Sc01, Sc02, Sc03, and Sc04 in sequence. Co-transformation of Y33-MBP-tTS-FoTO1 and Y22-ATR2-t24T5αOH into Sc-GGPP generated strain Sc05. Single colonies of strains Sc01–Sc05 were inoculated into test tubes containing 4 mL of tryptophan- and uracil-dropout CM medium and cultured overnight at 30 °C with shaking at 200 rpm. The seed cultures were then inoculated at a 1:50 ratio into 50 mL of tryptophan- and uracil-dropout CM medium in Erlenmeyer flasks and fermented at 30 °C and 200 rpm for 24 h. Subsequently, glucose and *n*-dodecane were added to final concentrations of 2% and 10%, respectively, and fermentation was continued for an additional 96 h. The fermentation broth was then centrifuged at 5,000 rpm for 10 min, and the upper organic phase was collected into glass vials for GC–MS analysis.

### Determination of taxadiene and its oxidation products by GC-MS

The products were detected by Agilent 7200 Accurate-Mass Quadrupole Time-of-Flight GC-MS System. The samples were injected into a TRACE DB-5MS column (30 m×0.25 mm×0.25 µm). The oven temperature was set at 80 °C for 1 min, increased to 220 °C at a rate of 10 °C/min, and held at 220 °C for 15 min. The injector and transfer lines were maintained at 230 °C and 240 °C, respectively. Mass detection was achieved with electric ionisation using SIM-scan mode with diagnostic ions of 288 (m/z).

Detailed information regarding strains, vectors, primers, and sequences is provided in Supplementary Tables 1-4.

## COMPUTATIONAL METHODS

### System construction and molecular dynamics simulations

The structure of FoTO1 was predicted using the Protenix^19^, and the model with the highest confidence score was selected for further analysis. Based on mechanistic requirements, the catalytic residues D68 and D149 were manually assigned to their protonated states (ASH). The geometry of the substrate, taxadiene-4(5)-epoxide, was optimized at the B3LYP-D4^21^/def2-SVP level of theory using the Orca^22^ software package, and RESP^23^ atomic charges were subsequently calculated using the Multiwfn^24^ program.

Substrate docking was executed through the Rosetta^25^. To ensure a productive binding orientation, a distance constraint of less than 2 Å was applied between the epoxide oxygen and the D68 proton. A total of 10000 poses were generated, from which the conformation with the most favorable binding energy score was selected as the starting point for molecular dynamics (MD) simulations.

Molecular dynamics (MD) simulations were performed using the Amber22^26^ package. The protein-substrate complex was solvated in an OPC^27^ water box with a minimum buffer distance of 10 Å between the protein atoms and the box edges. The system was neutralized by the addition of appropriate counterions. The ff19SB force field was utilized to describe the protein, while the parameters for the substrate were generated using the GAFF2 force field.

System refinement followed a two-stage energy minimization protocol employing combined steepest descent and conjugate gradient algorithms to first relax the side chains under backbone positional restraints and subsequently optimize the entire system without constraints. The complex was then heated from 0 K to 300 K over 50 ps in the NVT ensemble, followed by 50 ps of pressure equilibration at 1 atm and 300 K in the NPT ensemble to achieve density balance.

The production MD phase consisted of a 200 ns trajectory conducted under NPT conditions (1 atm, 300 K). To ensure adequate sampling of the reactive orientation within the active site, a harmonic distance restraint was applied between the epoxide oxygen of the substrate and the carboxyl proton of D68 during the initial 100 ns of the production run. This restraint was subsequently removed, allowing the system to evolve freely for the final 100 ns. This two-stage simulation strategy was employed to evaluate the intrinsic stability of the catalytic configuration and to confirm that the substrate maintains a near-attack conformation in the absence of external biasing potentials.

### Clustering analysis and representative snapshot selection

To characterize the stable conformational landscape of the FoTO1-substrate complex, molecular dynamics (MD) trajectories were analyzed upon system equilibration. The clustering analysis was restricted to the final 100 ns of the production run to focus exclusively on the equilibrated steady-state ensemble. The conformational space within this window was partitioned into five discrete clusters using the k-means algorithm based on the root-mean-square deviation (RMSD) of the substrate. The centroid structure from each of the five clusters was selected as a representative snapshot for subsequent electronic structure calculations. This five-cluster resolution ensured that the investigated configurations represent the thermodynamically accessible states of the enzyme-substrate complex.

### QM/MM calculations

Hybrid quantum mechanics/molecular mechanics (QM/MM) calculations were performed on the five representative snapshots identified via clustering to evaluate the reaction mechanism. The QM region included the taxadiene-4(5)-epoxide substrate and the D68-D149 catalytic dyad. To account for local protein flexibility, an active zone encompassing all atoms within 10 Å of the QM region was allowed to relax during geometry optimizations. All structural optimizations were executed at the B3LYP-D4/def2-SVP level of theory using the L-BFGS algorithm.

The localization of the first transition state (TS1) was achieved through an initial flexible scan along the reaction coordinate followed by refinement using the Dimer method^28^. The second transition state (TS2) was identified via a multi-resolution flexible scan, initiated at a resolution of 0.1 Å and subsequently refined to 0.02 Å. Each localized transition state was subjected to numerical frequency analysis to confirm the presence of a single imaginary frequency corresponding to the intended bond-breaking or bond-forming event. The Dimer method and related optimization protocols were implemented through the libdlfind Python interface of the DL-find^29^ library.

To ensure the precision of the energetic predictions, single-point energy calculations were performed at the refined B3LYP-D4/def2-TZVP level. The final potential energy surface (PES) reported in this study represents a Boltzmann-weighted average at 300 K, derived from the results of all five independent reaction pathways.

## CONTRIBUTIONS

H.J. conceived the idea and supervised the project. H.J., X.L., J.B. and J.L. designed the experiments. J.L. performed the chemical synthesis and all in vitro functional assays. X.L. conducted in vivo mutant functional experiments. Y.Z. assisted J.L. and X.L. with functional experiments. J.B. carried out all computational simulations, including structural modeling, complex structure prediction, and QM/MM calculations, and interpreted the catalytic mechanism.

H.C. and C.J. assisted with complex structure prediction. N.Z. and Y.L. provided constructive suggestions on the catalytic mechanism. J.B. and J.L. analyzed the data and wrote the manuscript with input from all authors. All authors reviewed and approved the final version of the manuscript.

‡These authors contributed equally.

## ACKNOWLEDGMENT

This work was supported by the National Natural Science Foundation of China (No. 32371499; 32571467), Natural Science Foundation of Wuhan (2025040601020158), Hubei Provincial Key R&D Major Projects (2025BCA006), Youth Science and Technology Talent Cultivation Special Program of Hubei Province (2025DJA024), Science and Technology Major Project of Guangxi (Guike AA24206048, AA24206050, JF2504850012), Hubei University of Technology High-Level Talent Research Startup Fund Program (4301/00960).

## Notes

### Competing Interest Statement

The authors have declared no competing interest.

