## Supporting Information for "FoTO1 is an epoxide isomerase in paclitaxel biosynthesis"

**Table S1 Primers used in this study**

| Name | Prime |
|---|---|
| linker-MBP-R | accaccaccaccaccaccagtctgcgcgtctttcag |
| FoTO1-linker-F | ggtggtggtggtggtggtatggctgagacaatgaatgaaaaagtagg |
| t10-FoTO1-R | tagcagccggatcttagcgcacgtgcagcatc |

| | |
|---|---|
| TAA-MBP-FoTO1-F | taagatccggctgctaacaaagc |
| t20-FoTO1-R | tagcagccggatcttatctgataccttcacgggcac |
| t24-FoTO1-R | tagcagccggatcttaacgggcacgacccac |
| t41-FoTO1-R | tagcagccggatcttagtgctgattaatgcggtcttcg |
| Y33-F | atttattggagaaagataacatatcatactttcc |
| ADH1+TDH3-R | tgtatatgagatagttgattgtatgcttggt |
| FoTO1(SC)-F | accaagcatacaatcaactatctcatatacaatggctgaaaccatgaacgaaaagg |
| FoTO1(SC)-R | agtatgatatgttatctttctccaataaattcatggtggggttgggtcct |
| FoTO1ΔC10AA-SC-R | agtatgatatgttatctttctccaataaattcatctgacgtgcaacatcaac |
| FoTO1ΔC20AA-SC-R | agtatgatatgttatctttctccaataaattcatctgataccttctctggctcta |
| FoTO1ΔC24AA-SC-R | agtatgatatgttatctttctccaataaattcatctggctctaccaaccaat |
| FoTO1ΔC41AA-SC-R | agtatgatatgttatctttctccaataaattcagtgttggttgattctgtcttcg |
| t24T5αOH-5F(SC) | actttttacaacaaatataaaaacaatgttctccattgctttgtccgc |
| t24T5αOH-3R(SC) | gacataactaattacatgagtcgactcatggccttgggaacaactt |
| ATR2-5F | cgaataaacacacataaacaaacaaaatgtcctcttcttcttcttcgtc |
| ATR2-3R | tcctcatcaagattgctttattctagattaccatacatctctaagatatcttc |
| PGK+TDH3-5F | tgtttttatatttgttgtaaaaagtagata |
| TDH3-5F(NE)-3R | tttgtttgtttatgtgtgtttattcg |
| Y22-3R(NE) | ataaagcaatcttgatgaggataatg |
| Y22-3R(NE) | tcatgtaattagttatgtcacgcttac |
| Y33-5F(NE) | atttattggagaaagataacatatca |
| Y33-3R(NE) | atgattgcaatgaaaagtttaagttaagc |
| tTS-F | ggtggtggtggtggtggtatgtcctcttccaccggtacctct |
| TS-R | taaacttttcattgcaatcatgaattccgattcagacctggattgggtcgatgt |
| MBP(SC)-F | gtttcgaataaacacacataaacaaacaaaatgtctaagattgaagaaggtaagttggtt atctg |

| | |
|---|---|
| MBP(SC)-R | accaccaccaccaccacccttagtaattctggtttgagcgtcc |
| ADH1+TDH3-F | tttgtttgtttatgtgtgtttattcgaaactaagt |
| ADH1+TDH3-R | tgtatatgagatagttgattgtatgcttggt |
| FoTO1(Y48F)-F | atttgatcaacatcttcacctgtattgccactccaag |
| FoTO1(Y48F)-R | gtggcaatacaggtgaagatgttgatcaaatgaggaataatgttacgg |
| FoTO1(Y48H)-F | atttgatcaacatccacacctgtattgccactcca |
| FoTO1(Y48H)-R | gtggcaatacaggtgtggatgttgatcaaatgaggaataatgt |
| FoTO1(Y48A)-F | atttgatcaacatcgctacctgtattgccactccaa |
| FoTO1(Y48A)-R | tggcaatacaggtagcgatgttgatcaaatgaggaat |
| FoTO1(Y60F)-F | gagatttggaaattttccatccagacgctacctttga |
| FoTO1(Y60F)-R | ggtagcgtctggatggaaaatttccaaatctcttggagtggc |
| FoTO1(Y60A)-F | gagatttggaaattgctcatccagacgctacctttga |
| FoTO1(Y60A)-R | ggtagcgtctggatgagcaatttccaaatctcttggagtggc |
| FoTO1(D68N)-F | acgctacctttgaaaacttcttcgtcagagctttcgg |
| FoTO1(D68N)-R | gctctgacgaagaagttttcaaaggtagcgtctggatgataaatt |
| FoTO1(D68A)-F | acgctacctttgaagctttcttcgtcagagctttcgg |
| FoTO1(D68A)-R | gctctgacgaagaaagcttcaaaggtagcgtctgga |
| FoTO1(D68R)-F | acgctacctttgaaagattcttcgtcagagctttcgg |
| FoTO1(D68R)-R | gctctgacgaagaatctttcaaaggtagcgtctgga |
| FoTO1(H83F)-F | aaatcaaatccattttctactctttcccaaagttggtctacga |
| FoTO1(H83F)-R | tttgggaaagagtagaaaatggatttgatttccttaataccgaaagct |
| FoTO1(H83A)-F | aaatcaaatccattgcttactctttcccaaagttggtct |
| FoTO1(H83A)-R | tttgggaaagagtaagcaatggatttgatttccttaataccgaa |
| FoTO1(H147F)-F | gtaagatcatcaagttcgaagacagaatcaaccaacaccc |
| FoTO1(H147F)-R | ttgattctgtcttcgaacttgatgatcttaccgttttcgatt |
| FoTO1(H147A)-F | gtaagatcatcaaggctgaagacagaatcaaccaacaccc |

| | |
|---|---|
| FoTO1(H147A)-R | ttgattctgtcttcagccttgatgatcttaccgttttcgatt |
| FoTO1(D149N)-F | atcatcaagcacgaaaacagaatcaaccaacacccagtta |
| FoTO1(D149N)-R | gtgttggttgattctgttttcgtgcttgatgatcttaccg |
| FoTO1(D149A)-F | gatcatcaagcacgaagctagaatcaaccaacacccagtt |
| FoTO1(D149A)-R | gtgttggttgattctagcttcgtgcttgatgatcttacc |
| FoTO1(D149H)-F | tcatcaagcacgaacacagaatcaaccaacacccag |
| FoTO1(D149H)-R | tgttggttgattctgtgttcgtgcttgatgatcttaccg |
| FoTO1(D149R)-F | atcatcaagcacgaaagaagaatcaaccaacacccagttag |
| FoTO1(D149R)-R | gtgttggttgattcttctttcgtgcttgatgatcttaccg |
| linker-MBP-R | accaccaccaccaccaccagtctgcgcgtctttcag |
| FoTO1-linker-F | ggtggtggtggtggtggtatggctgagacaatgaatgaaaaagtagg |
| t10-FoTO1-R | tagcagccggatcttagcgcacgtgcagcatc |
| TAA-MBP-FoTO1-F | taagatccggctgctaacaaagc |
| t20-FoTO1-R | tagcagccggatcttatctgataccttcacgggcac |
| t24-FoTO1-R | tagcagccggatcttaacgggcacgacccac |
| t41-FoTO1-R | tagcagccggatcttagtgctgattaatgcggtcttcg |
| Y33-F | atttattggagaaagataacatatcatactttcc |
| ADH1+TDH3-R | tgtatatgagatagttgattgtatgcttggt |
| FoTO1(SC)-F | accaagcatacaatcaactatctcatatacaatggctgaaaccatgaacgaaaagg |
| FoTO1(SC)-R | agtatgatatgttatctttctccaataaattcatggtggggttgggtcct |
| FoTO1ΔC10AA-SC-R | agtatgatatgttatctttctccaataaattcatctgacgtgcaacatcaac |
| FoTO1ΔC20AA-SC-R | agtatgatatgttatctttctccaataaattcatctgataccttctctggctcta |
| FoTO1ΔC24AA-SC-R | agtatgatatgttatctttctccaataaattcatctggctctaccaaccaat |
| FoTO1ΔC41AA-SC-R | agtatgatatgttatctttctccaataaattcagtgttggttgattctgtcttcg |

**Table S2 Plasmids used in this study**

| Name | Relevant characteristics | Source |
|---|---|---|
| pET28a-FoTO1 | pET28a vector, kanR, carrying FoTO1 gene | GenScript |
| Y33-MBP-tTS | YCplac33 vector, pTDH3-MBP-linker(GGGGGG)-tPFK1 | Stored in lab |
| Y22-ATR2-t24T5αOH | YCplac22 vector, pTDH3-ATR2-tTDH1, pPGK1-t24T5αOH-tCYC1 | Stored in lab |
| pET28a-MBP-FoTO1 | pET28a vector, kanR, carrying MBP-linker(GGGGGG)-FoTO1 gene | This study |
| pET28a-MBP-FoTO1-t10 | pET28a vector, kanR, carrying MBP-linker(GGGGGG)-FoTO1-t10 gene | This study |
| pET28a-MBP-FoTO1-t20 | pET28a vector, kanR, carrying MBP-linker(GGGGGG)-FoTO1-t20 gene | This study |
| pET28a-MBP-FoTO1-t24 | pET28a vector, kanR, carrying MBP-linker(GGGGGG)-FoTO1-t24 gene | This study |
| pET28a-MBP-FoTO1-t41 | pET28a vector, kanR, carrying MBP-linker(GGGGGG)-FoTO1-t41 gene | This study |
| Y33-MBP-tTS-FoTO1 | YCplac33 vector, pTDH3-MBP-linker(GGGGGG)-tPFK1, pADH1-FoTO1-tGPD1 | This study |
| Y33-MBP-tTS-FoTO1-t10 | YCplac33 vector, pTDH3-MBP-linker(GGGGGG)-tPFK1, pADH1- FoTO1-t10-tGPD1 | This study |
| Y33-MBP-tTS-FoTO1-t20 | YCplac33 vector, pTDH3-MBP-linker(GGGGGG)-tPFK1, pADH1- FoTO1-t20-tGPD1 | This study |
| Y33-MBP-tTS-FoTO1-t24 | YCplac33 vector, pTDH3-MBP-linker(GGGGGG)-tPFK1, pADH1- FoTO1-t24-tGPD1 | This study |
| Y33-MBP-tTS-FoTO1-t41 | YCplac33 vector, pTDH3-MBP-linker(GGGGGG)-tPFK1, pADH1- FoTO1-t41-tGPD1 | This study |

**Table S3 Strains used in this study.**

| Strain | Genotype | Reference |
|---|---|---|
| Sc-GGPP (BY4742-T2) | pPGK1-SaGGPPS-tADH1 and pPGK1-tHMG1-tADH1 cassettes integrated into δDNA site of BY4742 | [1] |
| Sc01 | Sc-GGPP carrying Y22-ATR2-t24T5αOH and Y33-MBP-tTS-FoTO1-t10 | This study |
| Sc02 | Sc-GGPP carrying Y22-ATR2-t24T5αOH and Y33-MBP-tTS-FoTO1-t20 | This study |
| Sc03 | Sc-GGPP carrying Y22-ATR2-t24T5αOH and Y33-MBP-tTS-FoTO1-t24 | This study |
| Sc04 | Sc-GGPP carrying Y22-ATR2-t24T5αOH and Y33-MBP-tTS-FoTO1-41 | This study |
| Sc05 | Sc-GGPP carrying Y22-ATR2-t24T5αOH and Y33-MBP-tTS-FoTO1 | This study |

**Table S4 Gene and amino acid sequences used in this study**

| Name | **Gene sequences** | **Amino acid sequences** |
|---|---|---|
| T5αOH (codon optimization for *Saccharomyces cerevisiae*) | atggatgctttgtacaagtccaccgttgccaagttcaacgaggttacccagttgga<br>ttgctccaccgaatccttctccattgctttgtccgctatcgccggcatcttgttgttgt<br>tgttgttgttcaggtccaagaggcattcctccttgaagttgccaccaggtaagttgg<br>gtatcccattcatcggcgagagcttcatcttcttgagggctttgaggtccaacagc<br>ttggaacagttcttcgacgagagagtcaagaagttcggcttggtcttcaagacctc<br>cttgatcggtcatccaaccgttgttctttgcggtccagcaggtaataggttgatctt<br>gtccaacgaggagaagttggtgcagatgtcttggccagcccaattcatgaagttg<br>atgggcgagaacagcgttgctactagaagaggcgaagaccatatcgtcatgag<br>gtcagctttggcaggtttctttggtccaggtgctttgcagtcctacatcggcaagat<br>gaacaccgagatccagtcccacatcaacgagaagtggaagggtaaggacgag | MDALYKSTVAKFNEVTQL<br>DCSTESFSIALSAIAGILLLL<br>LLFRSKRHSSLKLPPGKLGI<br>PFIGESFIFLRALRSNSLEQF<br>FDERVKKFGLVFKTSLIGHP<br>TVVLCGPAGNRLILSNEEKL<br>VQMSWPAQFMKLMGENSV<br>ATRRGEDHIVMRSALAGFF<br>GPGALQSYIGKMNTEIQSHI<br>NEKWKGKDEVNVLPLVRE |

| | | |
|---|---|---|
| | gtcaacgttttgccattggtcagggagttggtcttcaacatctccgccatcttgttct | LVFNISAILFFNIYDKQEQD |
| | tcaacatctacgacaagcaggagcaggacagattgcacaagttgttggagacc | RLHKLLETILVGSFALPIDLP |
| | atcttggtcggttctttcgctttgccaatcgacttgccaggtttcggtttccatagag | GFGFHRALQGRAKLNKIML |
| | ctttgcagggtagagccaagttgaacaagatcatgttgtccttgatcaagaagag | SLIKKRKEDLQSGSATATQ |
| | gaaggaagatttgcaatccggttccgctacagctacccaagacttgttgtccgtct | DLLSVLLTFRDDKGTPLTN |
| | tgttgaccttcagggacgataagggtactccattgaccaacgacgagatcttgga | DEILDNFSSLLHASYDTTTS |
| | caacttctcctccttgttgcacgcttcctacgatactaccacctctccaatggccttg | PMALIFKLLSSNPECYQKV |
| | atcttcaagttgttgtcctccaacccagagtgctaccaaaaggtcgttcaggagca | VQEQLEILSNKEEGEEITWK |
| | gttggagatcttgtccaacaaggaggaaggcgaagaaatcacctggaaggact | DLKAMKYTWQVAQETLR |
| | tgaaggccatgaagtacacttggcaagtcgctcaagagaccttgagaatgttccc | MFPPVFGTFRKAITDIQYDG |
| | accagtcttcggtaccttcagaaaggctatcaccgacatccaatacgacggttac | YTIPKGWKLLWTTYSTHPK |
| | accatcccaaagggttggaagttgctttggactacttactccacccacccaaagg | DLYFNEPEKFMPSRFDQEG |
| | acttgtacttcaacgagccagagaagttcatgccatccagattcgaccaagaag | KHVAPYTFLPFGGGQRSCV |
| | gtaagcacgtcgctccatacacatttttgccatttggcggcggtcaaagatcttgc | GWEFSKMEILLFVHHFVKT |
| | gtaggttgggagttctccaagatggagatcttgttgttcgtccaccacttcgtcaag | FSSYTPVDPDEKISGDPLPPL |
| | accttctcttcctacaccccagttgatccagacgaaaagatctcaggcgatccatt | PSKGFSIKLFPRP |
| | gccaccattgccatctaagggtttctccatcaagttgttcccaaggccatga | |
| ATR2 | atgtcctcttcttcttcttcgtcaacctccatgatcgatctcatggcagcaatcatca | MSSSSSSSTSMIDLMAAIIK |
| | aaggagagcctgtaattgtctccgacccagctaatgcctccgcttacgagtccgt | GEPVIVSDPANASAYESVA |
| | agctgctgaattatcctctatgcttatagagaatcgtcaattcgccatgattgttacc | AELSSMLIENRQFAMIVTTS |
| | acttccattgctgttcttattggttgcatcgttatgctcgtttggaggagatccggttc | IAVLIGCIVMLVWRRSGSG |
| | tgggaattcaaaacgtgtcgagcctcttaagcctttggttattaagcctcgtgagg | NSKRVEPLKPLVIKPREEEI |
| | aagagattgatgatgggcgtaagaaagttaccatctttttcggtacacaaactggt | DDGRKKVTIFFGTQTGTAE |
| | actgctgaaggttttgcaaaggctttaggagaagaagctaaagcaagatatgaa | GFAKALGEEAKARYEKTRF |
| | aagaccagattcaaaatcgttgatttggatgattacgcggctgatgatgatgagta | KIVDLDDYAADDDEYEEKL |
| | tgaggagaaattgaagaaagaggatgtggctttcttcttcttagccacatatggag | KKEDVAFFFLATYGDGEPT |
| | atggtgagcctaccgacaatgcagcgagattctacaaatggttcaccgaggggga | DNAARFYKWFTEGNDRGE |
| | atgacagaggagaatggcttaagaacttgaagtatggagtgtttggattaggaaa | WLKNLKYGVFGLGNRQYE |
| | cagacaatatgagcattttaataaggttgccaaagttgtagatgacattcttgtcga | HFNKVAKVVDDILVEQGA |
| | acaaggtgcacagcgtcttgtacaagttggtcttggagatgatgaccagtgtattg | QRLVQVGLGDDDQCIEDDF |
| | aagatgactttaccgcttggcgagaagcattgtggcccgagcttgatacaatact | TAWREALWPELDTILREEG |
| | gagggaagaaggggatacagctgttgccacaccatacactgcagctgtgttag | DTAVATPYTAAVLEYRVSI |
| | aatacagagtttctattcacgactctgaagatgccaaattcaatgatataaacatgg | HDSEDAKFNDINMANGNG |
| | caaatgggaatggttacactgtgtttgatgctcaacatccttacaaagcaaatgtc | YTVFDAQHPYKANVAVKR |
| | gctgttaaaagggagcttcatactcccgagtctgatcgttcttgtatccatttggaat | ELHTPESDRSCIHLEFDIAGS |
| | ttgacattgctggaagtggacttacgtatgaaactggagatcatgttggtgtacttt | GLTYETGDHVGVLCDNLSE |
| | gtgataacttaagtgaaactgtagatgaagctcttagattgctggatatgtcacctg | TVDEALRLLDMSPDTYFSL |
| | atacttatttctcacttcacgctgaaaaagaagacggcacaccaatcagcagctc | HAEKEDGTPISSSLPPPFPPC |
| | actgcctcctcccttcccaccttgcaacttgagaacagcgcttacacgatatgcat | NLRTALTRYACLLSSPKKS |

| | | |
|---|---|---|
| | gtcttttgagttctccaaagaagtctgctttagttgcgttggctgctcatgcatctgat<br>cctaccgaagcagaacgattaaaacaccttgcttcacctgctggaaaggttgatg<br>aatattcaaagtgggtagtagagagtcaaagaagtctacttgaggtgatggccga<br>gtttccttcagccaagccaccacttggtgtcttcttcgctggagttgctccaaggtt<br>gcagcctaggttctattcgatatcatcatcgcccaagattgctgaaactagaattca<br>cgtcacatgtgcactggtttatgagaaaatgccaactggcaggattcataaggga<br>gtgtgttccacttggatgaagaatgctgtgccttacgagaagagtgaaaactgttc<br>ctcggcgccgatatttgttaggcaatccaacttcaagcttccttctgattctaaggta<br>ccgatcatcatgatcggtccagggactggattagctccattcagaggattccttca<br>ggaaagactagcgttggtagaatctggtgttgaacttgggccatcagttttgttcttt<br>ggatgcagaaaccgtagaatggatttcatctacgaggaagagctccagcgattt<br>gttgagagtggtgctctcgcagagctaagtgtcgccttctctcgtgaaggaccca<br>ccaaagaatacgtacagcacaagatgatggacaaggcttctgatatctggaatat<br>gatctctcaaggagcttatttatatgtttgtggtgacgccaaaggcatggcaagag<br>atgttcacagatctctccacacaatagctcaagaacaggggtcaatggattcaac<br>taaagcagagggcttcgtgaagaatctgcaaacgagtggaagatatcttagaga<br>tgtatggtaa | ALVALAAHASDPTEAERLK<br>HLASPAGKVDEYSKWVVE<br>SQRSLLEVMAEFPSAKPPLG<br>VFFAGVAPRLQPRFYSISSS<br>PKIAETRIHVTCALVYEKMP<br>TGRIHKGVCSTWMKNAVP<br>YEKSENCSSAPIFVRQSNFK<br>LPSDSKVPIIMIGPGTGLAPF<br>RGFLQERLALVESGVELGP<br>SVLFFGCRNRRMDFIYEEEL<br>QRFVESGALAELSVAFSRE<br>GPTKEYVQHKMMDKASDI<br>WNMISQGAYLYVCGDAKG<br>MARDVHRSLHTIAQEQGS<br>MDSTKAEGFVKNLQTSGR<br>YLRDVW |
| FoTO1(codon optimization for *Saccharomyces cerevisiae*) | atggctgaaaccatgaacgaaaaggtcggtgctgatgaaaaaattgattgaagac<br>tcaagaaacactggccaagaagaaaacgactactccaagagattaggtccacta<br>tcccgtaacattattcctcatttgatcaacatctacacctgtattgccactccaagag<br>atttggaaatttatcatccagacgctacctttgaagatttcttcgtcagagctttcggt<br>attaaggaaatcaaatccattcactactctttcccaaagttggtctacgacggtaag<br>attttggaatactctgttgaagagaatgaaacttctccaggttgtggtgaattattgtt<br>caacatcaagcaacaatacaaggttttatacttgggtaaggaagttaacatcacta<br>ccttgatggttttgcaaatcgaaaacggtaagatcatcaagcacgaagacagaat<br>caaccaacacccagttagaggtagacacgatatttctgtcccattggttggtaga<br>gccagagaaggtatcagacgtttgactatgttgatgttgcacgtcagaatgggttt<br>cggtaaggacccaaccccaccatga | MAETMNEKVGADEKLIEDS<br>RNTGQEENDYSKRLGPLSR<br>NIIPHLINIYTCIATPRDLEIY<br>HPDATFEDFFVRAFGIKEIK<br>SIHYSFPKLVYDGKILEYSV<br>EENETSPGCGELLFNIKQQY<br>KVLYLGKEVNITTLMVLQI<br>ENGKIIKHEDRINQHPVRGR<br>HDISVPLVGRAREGIRRLTM<br>LMLHVRMGFGKDPTPP |
| MBP+Gly-linker+tTS(codon optimization for *Saccharomyces cerevisiae*) | atgtctaagattgaagaaggtaagttggttatctggattaacggtgacaagggtta<br>caacggtttggctgaagttggtaagaaatttgaaaaagataccggtatcaaggtc<br>actgttgaacacccagacaagttggaagaaaagtttccacaagttgctgccactg<br>gtgatggtccagacattatcttctgggctcatgacagattcggtggttacgcccaa<br>tccggtttgttagccgagatcaccccagataaggcttttcaagataagttgtatcca<br>ttcacttgggatgccgtcagatacaacggtaagttaatcgcctacccaattgctgtt<br>gaagctttgtctttgatctacaataaggacttgttacctaacccaccaaagacctgg<br>gaagaaatcccagctttagataaggagttaaaagctaagggtaagtccgctttga | MSKIEEGKLVIWINGDKGY<br>NGLAEVGKKFEKDTGIKVT<br>VEHPDKLEEKFPQVAATGD<br>GPDIIFWAHDRFGGYAQSG<br>LLAEITPDKAFQDKLYPFT<br>WDAVRYNGKLIAYPIAVEA<br>LSLIYNKDLLPNPPKTWEEI<br>PALDKELKAKGKSALMFNL |

| | |
|---|---|
| tgtttaacttgcaagaaccatacttcacttggccattgatcgctgctgatggtggtta | QEPYFTWPLIAADGGYAFK |
| cgcttttaagtatgaaaacggtaaatacgacattaaggatgtcggtgtcgacaatg | YENGKYDIKDVGVDNAGA |
| ctggtgctaaggccggtttaactttcttagtcgatttgattaagaataaacatatgaa | KAGLTFLVDLIKNKHMNA |
| tgctgacactgattactctattgctgaagctgctttcaacaagggtgaaaccgctat | DTDYSIAEAAFNKGETAMT |
| gactattaacggtccatgggcctggtctaacattgatacctctaaagtcaactacg | INGPWAWSNIDTSKVNYGV |
| gtgtcaccgtcttgccaacttttaagggtcaaccatctaagccattcgtcggtgtct | TVLPTFKGQPSKPFVGVLSA |
| tgtctgccggtattaacgctgcctctccaaataaggaattggccaaggaattctta | GINAASPNKELAKEFLENY |
| gaaaactacttgttaaccgatgaaggtttagaggccgttaacaaggataagccat | LLTDEGLEAVNKDKPLGAV |
| taggtgctgttgctttgaagtcttacgaagaagagttggctaaggatccaagaatt | ALKSYEEELAKDPRIAATM |
| gctgctactatggaaaacgctcaaaagggtgaaattatgccaaacatcccacaa | ENAQKGEIMPNIPQMSAFW |
| atgtctgctttctggtacgctgttcgtaccgccgtcattaatgccgcttctggtcgtc | YAVRTAVINAASGRQTVDE |
| aaactgttgatgaagccttgaaggacgctcaaaccagaattactaagggtggtg | ALKDAQTRITKggggggMSSS |
| gtggtggtggtatgtcctcttccaccggtacctctaaggttgtttccgaaacctcct | TGTSKVVSETSSTIVDDIPRL |
| ccaccattgttgacgacatcccaaggttgtccgctaattaccacggcgatctttgg | SANYHGDLWHHNVIQTLET |
| caccacaacgttatccaaaccttggagaccccattcagagaatcttccacctacc | PFRESSTYQERADELVVKIK |
| aagaaagagccgacgaattggtcgtcaagatcaaggacatgttcaacgccttgg | DMFNALGDGDISPSAYDTA |
| gtgacggagatatttctccatccgcttacgataccgcttgggttgctagagttgcta | WVARVATISSDGSEKPRFP |
| ctatctcctccgacggttcagaaaagccaagattcccacaggccttgaattgggt | QALNWVFNNQLQDGSWGI |
| cttcaacaaccagttgcaagacggttcttggggtatcgaatcccatttctccctttg | ESHFSLCDRLLNTTNSVIAL |
| cgacaggttgttgaacaccaccaactccgtcatcgctttgtccgtttggaagacc | SVWKTGHSQVEQGTEFIAE |
| ggtcattcccaagttgaacagggtaccgaattcatcgccgaaaacttgaggttgtt | NLRLLNEEDELSPDFEIIFPA |
| gaacgaggaggacgagttgtctccagatttcgagatcatcttcccagccttgttgc | LLQKAKALGINLPYDLPFIK |
| agaaggctaaggctttgggcatcaacttgccatacgacttgccattcatcaagtcc | SLSTTREARLTDVSAAADNI |
| ttgtccaccaccagagaagctagattgactgacgtttcagccgccgccgataata | PANMLNALEGLEEVIDWN |
| ttccagctaacatgttgaacgccttggaaggtttggaggaggtcatcgattggaa | KIMRFQSKDGSFLSSPASTA |
| caagatcatgagattccagtccaaggacggttccttcttgtcatctccagcttctac | CVLMNIGDEKCFTFLNNLL |
| cgcttgcgttttgatgaacatcggcgacgagaagtgcttcaccttcctaaacaact | DKFGGCVPCMYSIDLLERL |
| tgctagacaagttcggcggttgcgttccttgtatgtactctatcgacttgttggaga | SLVDNIEHLGIGRHFKQEIK |
| ggttgtccttggtcgataacatcgagcacttgggtatcggtaggcatttcaagcag | GALDYVYRHWSERGIGWG |
| gagatcaagggcgctttggattacgtctacaggcattggtccgaaagaggtatc | RDSLVPDLNTTALGLRTLR |
| ggttggggtagagactcattggttccagacttgaacaccaccgcattgggtttga | THGYDVSSDVLNNFKDEN |
| gaaccttgagaacccacggttacgacgtttcttccgacgtgttgaacaacttcaag | GRFFSSAGQTHVELRSVVN |
| gacgagaacggtagattcttctcctccgcaggtcaaacccacgttgaattgagat | LFRASDLAFPDEGVMDDAR |
| ccgtcgtcaacttgttcagggcttccgatttggcctttccagacgaaggagttatg | KFAEPYLRDALATKISTNTK |
| gacgacgctagaaagttcgccgaaccatacttgagagacgctttggctaccaag | LFKEIEYVVEYPWHMSIPRL |
| atctccaccaacaccaagttgttcaaggagatcgagtacgtcgtcgaataccctt | EAGSYIDSYDDDYVWQRK |
| ggcatatgtccatcccaaggttggaagccggttcttacatcgattcttacgacgac | TLYRMPSLSNSKCLELAKL |
| gactacgtttggcagaggaagaccttgtacaggatgccatccttgtccaacagca | DFNIVQSLHQEELKLLTRW |
| agtgtttggagctagccaagttggacttcaacatcgtgcaatccttgcaccagga | WKESGMADINFTRHRVAE |

| | |
|---|---|
| ggaattgaagttgctaaccaggtggtggaaggaatcaggtatggccgatatcaa | VYFSSATFEPEYSATRIAFT |
| cttcaccaggcacagagttgccgaagtttacttctcctccgctactttcgagccag | KIGCLQVLFDDMADIFATL |
| aatactccgctaccagaatcgctttcaccaagatcggttgcttgcaagtcttgttcg | DELKSFTEGVKRWDTSLLH |
| acgacatggccgatatcttcgctaccttggacgagttgaagtccttcaccgaagg | EIPECMQTCFKVWFKLMEE |
| agttaagaggtgggatacctccttgttgcacgaaatcccagagtgcatgcagact | VNNDVVKVQGRDMLAHIR |
| tgcttcaaggtttggttcaagttgatggaggaggtcaacaacgacgtcgtcaagg | KPWELYFNCYVQEREWLE |
| ttcaaggtagggatatgttggcccacatcagaaagccttgggagttgtacttcaat | AGYIPTFEEYLKTYAISVGL |
| tgctacgtccaggaaagggagtggttggaagcaggttacatcccaaccttcgag | GPCTLQPILLMGELVKDDV |
| gagtacctaaagacctacgctatctccgttggtttgggtccttgtactttgcagcca | VEKVHYPSNMFELVSLSWR |
| atcttgttgatgggcgagttggttaaggacgacgttgtcgaaaaggtccactacc | LTNDTKTYQAEKARGQQA |
| catccaacatgttcgagttggtgtccttgtcttggaggttgaccaacgacaccaag | SGIACYMKDNPGATEEDAI |
| acttaccaggccgaaaaggctagaggtcaacaagcttccggtatcgcttgctac | KHICRVVDRALKEASFEYF |
| atgaaggataacccaggagctacagaagaagacgctatcaagcacatttgcag | KPSNDIPMGCKSFIFNLRLC |
| ggtcgtcgacagagcattgaaggaagccagcttcgagtacttcaagccatctaa | VQIFYKFIDGYGIANEEIKD |
| cgacatcccaatgggttgcaagtccttcatcttcaacttgaggctttgcgtccagat | YIRKVYIDPIQV |
| cttctacaagttcatcgacggctacggtatcgctaacgaggagatcaaggactac | |
| atcaggaaggtctacatcgacccaatccaggtctga | |

**Table S5 Absolute QM/MM energies of stationary points.**

All values are reported in E (Hartree).

| | REA | TS1 | INT | TS2 | PRO |
|---|---|---|---|---|---|
| Snapshot 1 | -1314.5960 | -1314.5756 | -1314.5803 | -1314.5786 | -1314.6052 |
| Snapshot 2 | -1314.6029 | -1314.5840 | -1314.5890 | -1314.5858 | -1314.6167 |
| Snapshot 3 | -1314.5997 | -1314.5793 | -1314.5860 | -1314.5843 | -1314.6121 |
| Snapshot 4 | -1314.5971 | -1314.5748 | -1314.5806 | -1314.5783 | -1314.6143 |
| Snapshot 5 | -1314.5959 | -1314.5782 | -1314.5821 | -1314.5799 | -1314.6049 |

**Table S6 Geometric features of QM/MM optimized snapshots.**

Distances are reported in Å.

d(O-H): D68 proton to carboxyl oxygen.

d(H-OX): D68 proton to epoxide oxygen.

d(O-C): Epoxide oxygen to C4.

d(H-O): C20 methyl proton to D68 carboxyl oxygen.

| | | Snapshot 1 | Snapshot 2 | Snapshot 3 | Snapshot 4 | Snapshot 5 |
|---|---|---|---|---|---|---|
| DO-HOX | REA | 1.01 | 1.02 | 1.00 | 1.01 | 1.01 |
| | TS1 | 1.40 | 1.40 | 1.40 | 1.39 | 1.42 |
| | INT | 1.62 | 1.60 | 1.63 | 1.59 | 1.63 |
| | TS2 | 1.66 | 1.65 | 1.68 | 1.63 | 1.68 |
| | PRO | 1.87 | 1.73 | 1.85 | 1.74 | 1.88 |
| DOH-OX | REA | 1.61 | 1.55 | 1.64 | 1.61 | 1.63 |
| | TS1 | 1.07 | 1.07 | 1.07 | 1.07 | 1.07 |
| | INT | 1.00 | 1.00 | 1.00 | 1.00 | 1.00 |
| | TS2 | 0.99 | 0.99 | 0.99 | 0.99 | 0.97 |
| | PRO | 0.97 | 0.97 | 0.97 | 0.98 | 1.00 |
| XO-CX | REA | 1.47 | 1.47 | 1.47 | 1.47 | 1.47 |
| | TS1 | 1.91 | 1.91 | 1.95 | 1.90 | 1.90 |
| | INT | 2.31 | 2.31 | 2.33 | 2.31 | 2.32 |
| | TS2 | 2.33 | 2.34 | 2.35 | 2.33 | 2.34 |
| | PRO | 2.42 | 2.43 | 2.42 | 2.43 | 2.42 |
| XH-OD | REA | 2.65 | 2.63 | 2.64 | 2.75 | 2.77 |
| | TS1 | 2.09 | 2.09 | 2.08 | 2.00 | 2.13 |
| | INT | 1.81 | 1.81 | 1.80 | 1.74 | 1.83 |
| | TS2 | 1.50 | 1.46 | 1.48 | 1.48 | 1.50 |
| | PRO | 0.98 | 0.98 | 0.99 | 0.98 | 0.98 |

**Table S7 QM region coordinates of QM/MM optimized snapshots.**

All values are reported in Angstroms (Å).

| | | REA | TS1 | INT | TS2 | PRO |
|---|---|---|---|---|---|---|
| 1 | C | -0.751 0.997 -3.855 | -0.846 1.003 -3.661 | -0.774 1.046 -3.697 | -0.755 1.124 -3.726 | -0.769 1.111 -3.820 |
| | H | -0.826 0.335 -2.983 | -0.931 0.321 -2.804 | -0.759 0.362 -2.837 | -0.668 0.464 -2.853 | -0.626 0.461 -2.947 |
| | H | 0.261 1.429 -3.888 | 0.165 1.441 -3.662 | 0.208 1.539 -3.773 | 0.197 1.662 -3.859 | 0.138 1.717 -3.965 |
| | C | -1.742 2.134 -3.744 | -1.863 2.141 -3.546 | -1.850 2.111 -3.461 | -1.876 2.128 -3.440 | -1.947 2.006 -3.522 |
| | O | -1.749 3.103 -4.470 | -1.761 3.123 -4.287 | -1.881 3.117 -4.200 | -1.938 3.184 -4.132 | -2.068 3.053 -4.330 |
| | O | -2.568 1.979 -2.701 | -2.740 2.003 -2.603 | -2.609 1.924 -2.453 | -2.635 1.877 -2.466 | -2.698 1.822 -2.579 |
| | H | -3.127 2.790 -2.499 | -3.293 3.013 -1.798 | -3.171 2.888 -1.285 | -3.210 2.847 -1.247 | -3.383 2.873 -1.192 |

| | | | | | | | | | | | | | | | |
|---|---|---|---|---|---|---|---|---|---|---|---|---|---|---|---|
| C | -4.262 | -0.754 | -2.257 | -4.278 | -0.788 | -2.322 | -4.286 | -0.808 | -2.297 | -4.292 | -0.818 | -2.284 | -4.347 | -0.857 | -2.202 |
| H | -3.358 | -0.606 | -2.865 | -3.391 | -0.639 | -2.954 | -3.410 | -0.627 | -2.936 | -3.409 | -0.637 | -2.913 | -3.453 | -0.659 | -2.809 |
| H | -4.926 | 0.100 | -2.479 | -4.945 | 0.063 | -2.548 | -4.967 | 0.043 | -2.479 | -4.968 | 0.038 | -2.462 | -5.030 | -0.004 | -2.364 |
| C | -3.943 | -0.679 | -0.780 | -3.924 | -0.687 | -0.851 | -3.918 | -0.767 | -0.825 | -3.936 | -0.792 | -0.810 | -4.011 | -0.868 | -0.727 |
| O | -4.303 | -1.453 | 0.066 | -4.298 | -1.440 | 0.015 | -4.348 | -1.514 | 0.020 | -4.374 | -1.542 | 0.027 | -4.416 | -1.658 | 0.083 |
| O | -3.245 | 0.425 | -0.434 | -3.192 | 0.392 | -0.547 | -3.114 | 0.250 | -0.495 | -3.125 | 0.217 | -0.467 | -3.243 | 0.172 | -0.346 |
| H | -2.962 | 0.942 | -1.216 | -2.946 | 0.929 | -1.355 | -2.835 | 0.793 | -1.287 | -2.839 | 0.752 | -1.254 | -2.977 | 0.721 | -1.110 |
| C | 0.041 | 6.470 | -1.850 | 0.011 | 6.477 | -1.815 | 0.018 | 6.446 | -1.907 | 0.041 | 6.440 | -1.907 | 0.053 | 6.282 | -1.914 |
| C | -1.471 | 6.834 | -1.873 | -1.467 | 6.961 | -1.803 | -1.451 | 6.949 | -1.889 | -1.434 | 6.925 | -1.896 | -1.447 | 6.700 | -1.935 |
| C | -2.451 | 5.999 | -1.006 | -2.460 | 6.154 | -0.927 | -2.445 | 6.159 | -1.009 | -2.425 | 6.128 | -1.020 | -2.439 | 5.991 | -0.975 |
| C | -3.616 | 5.363 | -1.798 | -3.526 | 5.417 | -1.660 | -3.474 | 5.383 | -1.689 | -3.451 | 5.338 | -1.725 | -3.657 | 5.378 | -1.644 |
| O | -3.719 | 3.927 | -1.521 | -3.432 | 3.645 | -0.945 | -3.409 | 3.416 | -0.473 | -3.444 | 3.383 | -0.451 | -3.631 | 3.323 | -0.370 |
| C | -3.856 | 5.689 | -3.254 | -3.633 | 5.374 | -3.136 | -3.626 | 5.256 | -3.121 | -3.544 | 5.160 | -3.128 | -4.073 | 5.573 | -2.909 |
| C | -4.761 | 4.802 | -1.051 | -4.495 | 4.604 | -0.891 | -4.305 | 4.445 | -0.826 | -4.307 | 4.429 | -0.847 | -4.438 | 4.440 | -0.730 |
| C | -4.920 | 4.967 | 0.435 | -4.853 | 5.029 | 0.512 | -4.831 | 5.105 | 0.440 | -4.840 | 5.114 | 0.403 | -4.845 | 5.138 | 0.564 |
| C | -3.699 | 5.582 | 1.133 | -3.710 | 5.738 | 1.237 | -3.733 | 5.856 | 1.186 | -3.738 | 5.850 | 1.157 | -3.679 | 5.865 | 1.233 |
| C | -2.994 | 6.684 | 0.306 | -3.006 | 6.828 | 0.393 | -2.978 | 6.881 | 0.315 | -2.975 | 6.867 | 0.283 | -2.917 | 6.825 | 0.289 |
| C | -3.988 | 7.813 | -0.032 | -3.989 | 7.954 | 0.028 | -3.912 | 8.027 | -0.113 | -3.907 | 8.011 | -0.155 | -3.848 | 7.971 | -0.153 |
| C | -1.909 | 7.357 | 1.195 | -1.882 | 7.460 | 1.255 | -1.840 | 7.508 | 1.158 | -1.844 | 7.501 | 1.135 | -1.765 | 7.491 | 1.092 |
| C | -0.666 | 6.570 | 1.680 | -0.673 | 6.600 | 1.715 | -0.642 | 6.637 | 1.630 | -0.643 | 6.643 | 1.621 | -0.562 | 6.655 | 1.602 |
| C | 0.100 | 5.921 | 0.556 | 0.033 | 5.898 | 0.582 | 0.039 | 5.913 | 0.498 | 0.059 | 5.917 | 0.502 | 0.145 | 5.891 | 0.511 |
| C | 0.040 | 4.589 | 0.326 | -0.119 | 4.570 | 0.342 | -0.128 | 4.576 | 0.288 | -0.097 | 4.582 | 0.297 | 0.015 | 4.551 | 0.375 |
| C | 0.398 | 4.009 | -1.030 | 0.194 | 3.985 | -1.023 | 0.161 | 3.966 | -1.070 | 0.194 | 3.965 | -1.059 | 0.327 | 3.867 | -0.943 |
| C | 0.347 | 5.003 | -2.225 | 0.193 | 4.994 | -2.207 | 0.175 | 4.954 | -2.272 | 0.220 | 4.949 | -2.264 | 0.279 | 4.782 | -2.199 |
| C | -0.506 | 3.559 | 1.285 | -0.682 | 3.557 | 1.308 | -0.631 | 3.586 | 1.308 | -0.628 | 3.595 | 1.306 | -0.570 | 3.616 | 1.404 |
| C | 0.764 | 6.809 | -0.517 | 0.763 | 6.742 | -0.482 | 0.781 | 6.724 | -0.583 | 0.795 | 6.729 | -0.581 | 0.831 | 6.671 | -0.629 |
| C | 2.276 | 6.474 | -0.609 | 2.241 | 6.283 | -0.576 | 2.249 | 6.231 | -0.674 | 2.267 | 6.249 | -0.666 | 2.318 | 6.243 | -0.735 |
| C | 0.733 | 8.329 | -0.257 | 0.847 | 8.253 | -0.200 | 0.900 | 8.237 | -0.326 | 0.899 | 8.244 | -0.330 | 0.878 | 8.205 | -0.475 |
| H | -3.019 | 5.360 | -3.888 | -2.807 | 4.801 | -3.605 | -2.966 | 4.380 | -3.479 | -2.821 | 4.210 | -3.485 | -2.788 | 3.637 | -3.998 |
| H | 0.496 | 7.103 | -2.635 | 0.499 | 7.087 | -2.597 | 0.509 | 7.034 | -2.704 | 0.531 | 7.030 | -2.703 | 0.512 | 6.842 | -2.751 |
| H | -1.770 | 6.751 | -2.926 | -1.799 | 6.959 | -2.849 | -1.797 | 6.949 | -2.931 | -1.774 | 6.929 | -2.940 | -1.773 | 6.518 | -2.969 |
| H | -1.580 | 7.903 | -1.639 | -1.495 | 8.019 | -1.510 | -1.466 | 8.008 | -1.601 | -1.456 | 7.983 | -1.605 | -1.517 | 7.792 | -1.816 |
| H | -1.900 | 5.123 | -0.662 | -1.870 | 5.318 | -0.525 | -1.861 | 5.318 | -0.548 | -1.850 | 5.296 | -0.559 | -1.934 | 5.123 | -0.548 |
| H | -4.015 | 6.769 | -3.390 | -3.586 | 6.404 | -3.526 | -3.290 | 6.119 | -3.707 | -3.152 | 5.966 | -3.759 | -3.591 | 6.277 | -3.591 |
| H | -4.762 | 5.172 | -3.603 | -4.575 | 4.910 | -3.453 | -4.643 | 4.939 | -3.395 | -4.522 | 4.782 | -3.457 | -4.996 | 5.105 | -3.263 |
| H | -5.705 | 4.689 | -1.596 | -5.382 | 4.289 | -1.456 | -5.152 | 4.053 | -1.412 | -5.155 | 4.041 | -1.434 | -5.353 | 4.099 | -1.246 |
| H | -5.178 | 3.983 | 0.867 | -5.177 | 4.128 | 1.060 | -5.241 | 4.290 | 1.058 | -5.275 | 4.315 | 1.025 | -5.240 | 4.355 | 1.230 |
| H | -5.803 | 5.613 | 0.573 | -5.736 | 5.685 | 0.433 | -5.667 | 5.774 | 0.181 | -5.660 | 5.794 | 0.126 | -5.671 | 5.833 | 0.347 |
| H | -2.965 | 4.799 | 1.369 | -2.965 | 4.995 | 1.554 | -3.015 | 5.129 | 1.594 | -3.030 | 5.116 | 1.567 | -2.975 | 5.124 | 1.640 |

| | | | | | | | | | | | | | | | | |
|---|---|---|---|---|---|---|---|---|---|---|---|---|---|---|---|---|
| | H | -4.016 | 6.006 | 2.100 | -4.095 | 6.201 | 2.159 | -4.168 | 6.389 | 2.048 | -4.166 | 6.385 | 2.020 | -4.049 | 6.439 | 2.099 |
| | H | -4.810 | 7.489 | -0.684 | -4.824 | 7.598 | -0.596 | -4.758 | 7.668 | -0.717 | -4.747 | 7.653 | -0.768 | -4.735 | 7.617 | -0.697 |
| | H | -4.436 | 8.217 | 0.891 | -4.419 | 8.410 | 0.935 | -4.321 | 8.543 | 0.770 | -4.324 | 8.528 | 0.725 | -4.198 | 8.546 | 0.720 |
| | H | -3.484 | 8.650 | -0.540 | -3.486 | 8.755 | -0.537 | -3.386 | 8.779 | -0.719 | -3.371 | 8.763 | -0.754 | -3.328 | 8.680 | -0.816 |
| | H | -2.431 | 7.717 | 2.096 | -2.371 | 7.834 | 2.171 | -2.322 | 7.905 | 2.068 | -2.334 | 7.897 | 2.040 | -2.231 | 7.942 | 1.985 |
| | H | -1.587 | 8.273 | 0.690 | -1.525 | 8.360 | 0.742 | -1.468 | 8.395 | 0.631 | -1.475 | 8.389 | 0.609 | -1.402 | 8.345 | 0.510 |
| | H | -0.987 | 5.823 | 2.413 | -1.015 | 5.874 | 2.460 | -0.991 | 5.925 | 2.385 | -0.993 | 5.932 | 2.377 | -0.912 | 5.971 | 2.384 |
| | H | -0.040 | 7.281 | 2.245 | 0.009 | 7.278 | 2.255 | 0.052 | 7.311 | 2.157 | 0.042 | 7.326 | 2.149 | 0.124 | 7.357 | 2.108 |
| | H | -0.315 | 3.184 | -1.211 | -0.560 | 3.203 | -1.208 | -0.612 | 3.195 | -1.233 | -0.586 | 3.202 | -1.219 | -0.409 | 3.052 | -1.052 |
| | H | 1.387 | 3.516 | -0.988 | 1.156 | 3.441 | -0.988 | 1.113 | 3.405 | -1.025 | 1.138 | 3.391 | -1.017 | 1.306 | 3.358 | -0.883 |
| | H | 1.325 | 4.985 | -2.729 | 1.157 | 4.905 | -2.726 | 1.136 | 4.848 | -2.793 | 1.193 | 4.847 | -2.767 | 1.235 | 4.679 | -2.734 |
| | H | -0.376 | 4.650 | -2.976 | -0.561 | 4.695 | -2.948 | -0.585 | 4.655 | -3.007 | -0.526 | 4.645 | -3.010 | -0.495 | 4.430 | -2.896 |
| | H | 0.155 | 2.675 | 1.318 | 0.008 | 2.698 | 1.381 | 0.049 | 2.716 | 1.327 | 0.047 | 2.723 | 1.353 | 0.057 | 2.712 | 1.488 |
| | H | -0.608 | 3.918 | 2.316 | -0.820 | 3.942 | 2.325 | -0.667 | 3.982 | 2.330 | -0.695 | 4.002 | 2.322 | -0.636 | 4.053 | 2.407 |
| | H | -1.493 | 3.184 | 0.953 | -1.642 | 3.149 | 0.947 | -1.626 | 3.196 | 1.038 | -1.619 | 3.212 | 1.011 | -1.579 | 3.276 | 1.107 |
| | H | 2.732 | 7.011 | -1.461 | 2.746 | 6.796 | -1.413 | 2.759 | 6.706 | -1.532 | 2.777 | 6.733 | -1.517 | 2.780 | 6.696 | -1.630 |
| | H | 2.791 | 6.801 | 0.308 | 2.776 | 6.545 | 0.350 | 2.797 | 6.513 | 0.239 | 2.808 | 6.528 | 0.252 | 2.878 | 6.593 | 0.146 |
| | H | 2.485 | 5.404 | -0.733 | 2.356 | 5.203 | -0.724 | 2.342 | 5.145 | -0.786 | 2.369 | 5.163 | -0.785 | 2.459 | 5.157 | -0.796 |
| | H | 1.070 | 8.565 | 0.763 | 1.229 | 8.447 | 0.813 | 1.299 | 8.439 | 0.677 | 1.282 | 8.450 | 0.680 | 1.193 | 8.500 | 0.537 |
| | H | 1.423 | 8.822 | -0.960 | 1.549 | 8.707 | -0.917 | 1.604 | 8.666 | -1.056 | 1.611 | 8.674 | -1.052 | 1.615 | 8.613 | -1.186 |
| | H | -0.241 | 8.804 | -0.409 | -0.093 | 8.801 | -0.313 | -0.030 | 8.802 | -0.437 | -0.034 | 8.802 | -0.453 | -0.067 | 8.713 | -0.693 |
| 2 | C | -2.018 | 4.197 | -2.534 | -2.034 | 4.217 | -2.418 | -1.994 | 4.301 | -2.422 | -1.985 | 4.388 | -2.413 | -2.099 | 4.531 | -2.653 |
| | H | -2.061 | 3.247 | -1.985 | -2.040 | 3.264 | -1.871 | -1.928 | 3.357 | -1.863 | -1.860 | 3.457 | -1.843 | -2.088 | 3.641 | -2.012 |
| | H | -0.981 | 4.573 | -2.517 | -1.016 | 4.636 | -2.392 | -1.001 | 4.779 | -2.432 | -1.022 | 4.923 | -2.443 | -1.123 | 5.038 | -2.619 |
| | C | -2.904 | 5.246 | -1.898 | -2.975 | 5.232 | -1.770 | -2.987 | 5.238 | -1.730 | -3.024 | 5.249 | -1.695 | -3.209 | 5.434 | -2.174 |
| | O | -2.931 | 6.407 | -2.241 | -2.943 | 6.407 | -2.142 | -3.042 | 6.432 | -2.087 | -3.112 | 6.476 | -1.993 | -3.075 | 6.718 | -2.525 |
| | O | -3.613 | 4.757 | -0.872 | -3.715 | 4.766 | -0.812 | -3.663 | 4.728 | -0.777 | -3.701 | 4.694 | -0.792 | -4.141 | 5.038 | -1.508 |
| | H | -4.090 | 5.466 | -0.321 | -4.214 | 5.480 | 0.284 | -4.123 | 5.295 | 0.648 | -4.138 | 5.219 | 0.707 | -4.176 | 5.122 | 0.221 |
| | C | -5.185 | 1.934 | -1.283 | -5.177 | 1.973 | -1.320 | -5.182 | 1.972 | -1.319 | -5.189 | 1.956 | -1.306 | -5.216 | 2.000 | -1.002 |
| | H | -4.385 | 2.326 | -1.927 | -4.395 | 2.365 | -1.986 | -4.409 | 2.379 | -1.988 | -4.417 | 2.368 | -1.971 | -4.369 | 2.418 | -1.567 |
| | H | -5.882 | 2.776 | -1.125 | -5.872 | 2.815 | -1.159 | -5.884 | 2.805 | -1.139 | -5.891 | 2.786 | -1.115 | -5.895 | 2.853 | -0.830 |
| | C | -4.660 | 1.533 | 0.079 | -4.609 | 1.598 | 0.033 | -4.601 | 1.580 | 0.024 | -4.605 | 1.546 | 0.030 | -4.740 | 1.459 | 0.330 |
| | O | -4.906 | 0.505 | 0.654 | -4.855 | 0.579 | 0.633 | -4.869 | 0.568 | 0.626 | -4.881 | 0.534 | 0.627 | -4.723 | 0.293 | 0.633 |
| | O | -3.907 | 2.478 | 0.677 | -3.829 | 2.539 | 0.579 | -3.782 | 2.495 | 0.553 | -3.774 | 2.450 | 0.564 | -4.283 | 2.371 | 1.209 |
| | H | -3.761 | 3.263 | 0.103 | -3.720 | 3.343 | -0.011 | -3.667 | 3.301 | -0.035 | -3.662 | 3.249 | -0.020 | -4.490 | 3.329 | 1.061 |
| | C | -1.091 | 9.107 | 1.014 | -1.175 | 9.114 | 0.975 | -1.166 | 9.125 | 0.977 | -1.141 | 9.121 | 0.966 | -1.139 | 8.850 | 0.882 |
| | C | -2.595 | 9.414 | 1.277 | -2.678 | 9.442 | 1.212 | -2.661 | 9.480 | 1.215 | -2.642 | 9.453 | 1.192 | -2.666 | 9.052 | 1.101 |
| | C | -3.448 | 8.273 | 1.894 | -3.537 | 8.354 | 1.907 | -3.544 | 8.394 | 1.872 | -3.516 | 8.358 | 1.845 | -3.487 | 7.910 | 1.749 |
| | C | -4.609 | 7.759 | 1.019 | -4.607 | 7.735 | 1.074 | -4.574 | 7.771 | 1.046 | -4.545 | 7.723 | 0.996 | -4.702 | 7.460 | 0.955 |

| | | | | | | | | | | | | | | | |
|---|---|---|---|---|---|---|---|---|---|---|---|---|---|---|---|
| O | -4.544 | 6.290 | 0.914 | -4.315 | 5.861 | 1.283 | -4.305 | 5.547 | 1.596 | -4.321 | 5.485 | 1.638 | -4.373 | 5.065 | 1.173 |
| C | -4.966 | 8.432 | -0.281 | -4.841 | 8.077 | -0.343 | -4.808 | 8.050 | -0.349 | -4.735 | 7.951 | -0.382 | -5.275 | 8.116 | -0.067 |
| C | -5.636 | 6.892 | 1.637 | -5.445 | 6.663 | 1.659 | -5.288 | 6.562 | 1.630 | -5.273 | 6.531 | 1.615 | -5.300 | 6.144 | 1.437 |
| C | -5.706 | 6.661 | 3.122 | -5.711 | 6.682 | 3.145 | -5.738 | 6.759 | 3.068 | -5.741 | 6.773 | 3.041 | -5.603 | 6.195 | 2.931 |
| C | -4.474 | 7.148 | 3.895 | -4.578 | 7.307 | 3.962 | -4.635 | 7.355 | 3.935 | -4.637 | 7.365 | 3.910 | -4.466 | 6.781 | 3.776 |
| C | -3.905 | 8.490 | 3.378 | -4.022 | 8.629 | 3.381 | -4.031 | 8.657 | 3.370 | -4.018 | 8.651 | 3.327 | -3.936 | 8.138 | 3.251 |
| C | -4.976 | 9.598 | 3.452 | -5.098 | 9.729 | 3.376 | -5.074 | 9.789 | 3.354 | -5.058 | 9.787 | 3.290 | -5.046 | 9.208 | 3.320 |
| C | -2.759 | 8.947 | 4.320 | -2.872 | 9.115 | 4.301 | -2.881 | 9.112 | 4.300 | -2.875 | 9.117 | 4.266 | -2.812 | 8.654 | 4.190 |
| C | -1.443 | 8.135 | 4.430 | -1.569 | 8.277 | 4.434 | -1.593 | 8.253 | 4.433 | -1.582 | 8.269 | 4.421 | -1.457 | 7.912 | 4.315 |
| C | -0.767 | 7.867 | 3.113 | -0.908 | 7.948 | 3.122 | -0.946 | 7.933 | 3.114 | -0.908 | 7.946 | 3.114 | -0.771 | 7.663 | 3.000 |
| C | -0.831 | 6.657 | 2.509 | -0.990 | 6.717 | 2.552 | -1.052 | 6.706 | 2.529 | -0.990 | 6.715 | 2.543 | -0.772 | 6.444 | 2.412 |
| C | -0.624 | 6.523 | 1.015 | -0.767 | 6.528 | 1.064 | -0.829 | 6.527 | 1.042 | -0.777 | 6.529 | 1.054 | -0.545 | 6.298 | 0.920 |
| C | -0.848 | 7.822 | 0.191 | -0.921 | 7.806 | 0.196 | -0.933 | 7.820 | 0.187 | -0.888 | 7.813 | 0.189 | -0.799 | 7.579 | 0.078 |
| C | -1.252 | 5.362 | 3.154 | -1.404 | 5.444 | 3.244 | -1.438 | 5.428 | 3.227 | -1.387 | 5.441 | 3.241 | -1.130 | 5.139 | 3.081 |
| C | -0.211 | 9.047 | 2.296 | -0.320 | 9.090 | 2.273 | -0.328 | 9.074 | 2.286 | -0.304 | 9.086 | 2.275 | -0.276 | 8.864 | 2.174 |
| C | 1.274 | 8.772 | 1.943 | 1.166 | 8.777 | 1.964 | 1.155 | 8.731 | 1.995 | 1.183 | 8.760 | 1.983 | 1.226 | 8.673 | 1.849 |
| C | -0.184 | 10.412 | 3.012 | -0.291 | 10.482 | 2.934 | -0.277 | 10.460 | 2.962 | -0.269 | 10.475 | 2.942 | -0.348 | 10.242 | 2.861 |
| H | -5.822 | 7.914 | -0.735 | -5.728 | 7.568 | -0.739 | -5.808 | 7.725 | -0.667 | -5.695 | 7.579 | -0.764 | -6.168 | 7.708 | -0.547 |
| H | -0.730 | 9.957 | 0.409 | -0.805 | 9.945 | 0.349 | -0.775 | 9.956 | 0.363 | -0.756 | 9.951 | 0.348 | -0.828 | 9.714 | 0.267 |
| H | -3.022 | 9.719 | 0.312 | -3.101 | 9.685 | 0.229 | -3.077 | 9.760 | 0.239 | -3.056 | 9.729 | 0.213 | -3.068 | 9.255 | 0.099 |
| H | -2.671 | 10.319 | 1.897 | -2.751 | 10.382 | 1.777 | -2.718 | 10.403 | 1.806 | -2.713 | 10.377 | 1.781 | -2.832 | 9.996 | 1.641 |
| H | -2.804 | 7.398 | 1.932 | -2.862 | 7.499 | 2.045 | -2.884 | 7.504 | 2.032 | -2.866 | 7.477 | 2.012 | -2.857 | 7.017 | 1.780 |
| H | -4.139 | 8.394 | -1.002 | -3.991 | 7.807 | -0.994 | -4.104 | 7.379 | -0.972 | -3.959 | 7.198 | -1.048 | -3.805 | 7.224 | -2.105 |
| H | -5.242 | 9.485 | -0.114 | -4.981 | 9.169 | -0.413 | -4.579 | 9.073 | -0.671 | -4.440 | 8.930 | -0.779 | -4.941 | 9.101 | -0.400 |
| H | -6.598 | 6.792 | 1.118 | -6.350 | 6.411 | 1.087 | -6.158 | 6.298 | 1.005 | -6.139 | 6.261 | 0.988 | -6.240 | 5.940 | 0.899 |
| H | -5.889 | 5.586 | 3.302 | -5.899 | 5.641 | 3.456 | -6.018 | 5.755 | 3.423 | -6.043 | 5.782 | 3.416 | -5.860 | 5.173 | 3.259 |
| H | -6.610 | 7.193 | 3.460 | -6.655 | 7.234 | 3.291 | -6.646 | 7.379 | 3.090 | -6.639 | 7.410 | 3.038 | -6.512 | 6.807 | 3.045 |
| H | -3.684 | 6.384 | 3.842 | -3.754 | 6.584 | 4.048 | -3.836 | 6.610 | 4.062 | -3.845 | 6.612 | 4.045 | -3.632 | 6.065 | 3.818 |
| H | -4.734 | 7.251 | 4.962 | -4.934 | 7.494 | 4.987 | -5.028 | 7.562 | 4.944 | -5.030 | 7.588 | 4.915 | -4.816 | 6.908 | 4.815 |
| H | -5.340 | 9.720 | 4.486 | -5.468 | 9.921 | 4.396 | -5.436 | 9.992 | 4.376 | -5.438 | 9.992 | 4.304 | -5.343 | 9.377 | 4.368 |
| H | -4.557 | 10.564 | 3.130 | -4.696 | 10.676 | 2.983 | -4.647 | 10.723 | 2.955 | -4.620 | 10.721 | 2.905 | -4.695 | 10.170 | 2.914 |
| H | -5.846 | 9.399 | 2.809 | -5.961 | 9.466 | 2.747 | -5.941 | 9.543 | 2.723 | -5.916 | 9.547 | 2.646 | -5.943 | 8.931 | 2.752 |
| H | -3.189 | 8.984 | 5.335 | -3.302 | 9.194 | 5.314 | -3.323 | 9.178 | 5.309 | -3.325 | 9.192 | 5.271 | -3.249 | 8.682 | 5.203 |
| H | -2.531 | 9.992 | 4.080 | -2.625 | 10.146 | 4.022 | -2.614 | 10.143 | 4.045 | -2.612 | 10.148 | 4.003 | -2.636 | 9.707 | 3.943 |
| H | -1.657 | 7.187 | 4.928 | -1.797 | 7.353 | 4.971 | -1.838 | 7.325 | 4.955 | -1.828 | 7.342 | 4.944 | -1.628 | 6.960 | 4.821 |
| H | -0.784 | 8.694 | 5.114 | -0.895 | 8.846 | 5.095 | -0.909 | 8.803 | 5.099 | -0.915 | 8.829 | 5.097 | -0.831 | 8.511 | 4.997 |
| H | -1.335 | 5.752 | 0.671 | -1.500 | 5.772 | 0.735 | -1.588 | 5.800 | 0.704 | -1.540 | 5.798 | 0.733 | -1.227 | 5.496 | 0.584 |
| H | 0.372 | 6.096 | 0.797 | 0.217 | 6.058 | 0.882 | 0.141 | 6.025 | 0.862 | 0.187 | 6.022 | 0.865 | 0.470 | 5.906 | 0.721 |
| H | 0.050 | 7.982 | -0.419 | 0.013 | 7.940 | -0.365 | 0.013 | 7.938 | -0.358 | 0.059 | 7.934 | -0.354 | 0.108 | 7.789 | -0.504 |

| | | | | | | | | | | | | | | | | |
|---|---|---|---|---|---|---|---|---|---|---|---|---|---|---|---|---|
| | H | -1.675 | 7.669 | -0.517 | -1.702 | 7.644 | -0.559 | -1.708 | 7.689 | -0.581 | -1.658 | 7.673 | -0.582 | -1.594 | 7.390 | -0.658 |
| | H | -1.363 | 5.419 | 4.241 | -1.482 | 5.537 | 4.331 | -1.484 | 5.521 | 4.317 | -1.423 | 5.535 | 4.330 | -1.174 | 5.197 | 4.175 |
| | H | -2.202 | 4.992 | 2.724 | -2.365 | 5.067 | 2.849 | -2.404 | 5.045 | 2.856 | -2.364 | 5.074 | 2.884 | -2.099 | 4.756 | 2.711 |
| | H | -0.497 | 4.583 | 2.943 | -0.655 | 4.658 | 3.040 | -0.685 | 4.654 | 3.000 | -0.646 | 4.655 | 3.010 | -0.376 | 4.374 | 2.827 |
| | H | 1.876 | 8.771 | 2.866 | 1.752 | 8.813 | 2.896 | 1.729 | 8.741 | 2.935 | 1.759 | 8.790 | 2.921 | 1.816 | 8.741 | 2.777 |
| | H | 1.440 | 7.802 | 1.459 | 1.327 | 7.783 | 1.529 | 1.299 | 7.740 | 1.550 | 1.339 | 7.765 | 1.550 | 1.461 | 7.703 | 1.397 |
| | H | 1.672 | 9.564 | 1.285 | 1.586 | 9.530 | 1.276 | 1.604 | 9.481 | 1.323 | 1.618 | 9.509 | 1.300 | 1.582 | 9.466 | 1.171 |
| | H | -1.168 | 10.841 | 3.222 | -1.273 | 10.939 | 3.085 | -1.252 | 10.931 | 3.114 | -1.250 | 10.939 | 3.086 | -1.363 | 10.621 | 3.006 |
| | H | 0.368 | 10.354 | 3.962 | 0.220 | 10.455 | 3.909 | 0.229 | 10.414 | 3.939 | 0.231 | 10.440 | 3.921 | 0.153 | 10.231 | 3.842 |
| | H | 0.339 | 11.139 | 2.371 | 0.275 | 11.169 | 2.286 | 0.301 | 11.145 | 2.322 | 0.307 | 11.161 | 2.302 | 0.172 | 10.981 | 2.232 |
| 3 | C | -0.224 | -2.455 | -4.423 | -0.238 | -2.566 | -4.326 | -0.188 | -2.395 | -4.362 | -0.182 | -2.315 | -4.340 | -0.154 | -2.254 | -4.369 |
| | H | 0.039 | -3.380 | -3.892 | 0.024 | -3.503 | -3.816 | 0.204 | -3.302 | -3.881 | 0.282 | -3.193 | -3.872 | 0.377 | -3.107 | -3.927 |
| | H | 0.703 | -1.926 | -4.692 | 0.692 | -2.041 | -4.597 | 0.661 | -1.764 | -4.671 | 0.613 | -1.625 | -4.661 | 0.578 | -1.497 | -4.688 |
| | C | -1.035 | -1.544 | -3.526 | -1.044 | -1.669 | -3.384 | -1.039 | -1.631 | -3.339 | -1.052 | -1.638 | -3.276 | -1.046 | -1.695 | -3.286 |
| | O | -1.273 | -0.386 | -3.777 | -1.187 | -0.473 | -3.653 | -1.313 | -0.434 | -3.563 | -1.356 | -0.422 | -3.441 | -1.450 | -0.452 | -3.480 |
| | O | -1.405 | -2.143 | -2.383 | -1.477 | -2.247 | -2.308 | -1.352 | -2.265 | -2.277 | -1.356 | -2.314 | -2.257 | -1.328 | -2.332 | -2.283 |
| | H | -1.852 | -1.514 | -1.740 | -1.749 | -1.597 | -1.102 | -1.621 | -1.801 | -0.737 | -1.621 | -1.862 | -0.656 | -1.652 | -1.885 | -0.522 |
| | C | -3.045 | -4.803 | -1.984 | -3.157 | -4.848 | -2.051 | -3.149 | -4.803 | -2.032 | -3.161 | -4.819 | -2.004 | -3.178 | -4.844 | -1.952 |
| | H | -2.436 | -4.394 | -2.802 | -2.619 | -4.376 | -2.883 | -2.632 | -4.301 | -2.861 | -2.630 | -4.321 | -2.825 | -2.615 | -4.358 | -2.758 |
| | H | -3.666 | -3.962 | -1.629 | -3.806 | -4.055 | -1.638 | -3.794 | -4.032 | -1.574 | -3.804 | -4.046 | -1.547 | -3.816 | -4.063 | -1.500 |
| | C | -2.196 | -5.215 | -0.799 | -2.222 | -5.242 | -0.924 | -2.191 | -5.239 | -0.941 | -2.221 | -5.279 | -0.906 | -2.281 | -5.349 | -0.838 |
| | O | -2.357 | -6.196 | -0.118 | -2.368 | -6.196 | -0.194 | -2.373 | -6.175 | -0.194 | -2.428 | -6.212 | -0.165 | -2.531 | -6.287 | -0.122 |
| | O | -1.249 | -4.316 | -0.469 | -1.234 | -4.370 | -0.713 | -1.143 | -4.436 | -0.784 | -1.152 | -4.502 | -0.734 | -1.187 | -4.606 | -0.623 |
| | H | -1.200 | -3.582 | -1.117 | -1.229 | -3.637 | -1.392 | -1.128 | -3.696 | -1.457 | -1.114 | -3.772 | -1.405 | -1.107 | -3.882 | -1.277 |
| | C | 1.395 | 1.942 | -0.950 | 1.383 | 2.024 | -0.938 | 1.371 | 2.026 | -0.948 | 1.387 | 2.011 | -0.954 | 1.419 | 1.938 | -1.000 |
| | C | 0.004 | 2.177 | -0.290 | 0.005 | 2.310 | -0.274 | 0.009 | 2.331 | -0.263 | 0.013 | 2.300 | -0.289 | 0.006 | 2.165 | -0.394 |
| | C | -0.650 | 1.006 | 0.488 | -0.650 | 1.148 | 0.517 | -0.668 | 1.177 | 0.514 | -0.664 | 1.146 | 0.488 | -0.658 | 1.039 | 0.444 |
| | C | -2.048 | 0.586 | -0.019 | -1.919 | 0.613 | -0.035 | -1.896 | 0.624 | -0.037 | -1.901 | 0.586 | -0.085 | -2.022 | 0.577 | -0.037 |
| | O | -2.110 | -0.861 | -0.254 | -1.616 | -1.317 | -0.074 | -1.611 | -1.667 | 0.253 | -1.632 | -1.713 | 0.320 | -1.726 | -1.782 | 0.438 |
| | C | -2.765 | 1.381 | -1.085 | -2.531 | 1.087 | -1.294 | -2.498 | 1.019 | -1.289 | -2.447 | 0.921 | -1.345 | -2.750 | 1.121 | -1.032 |
| | C | -2.876 | -0.301 | 0.827 | -2.572 | -0.535 | 0.640 | -2.475 | -0.618 | 0.628 | -2.490 | -0.633 | 0.623 | -2.557 | -0.655 | 0.688 |
| | C | -2.488 | -0.672 | 2.235 | -2.439 | -0.656 | 2.141 | -2.506 | -0.525 | 2.146 | -2.552 | -0.481 | 2.136 | -2.584 | -0.442 | 2.197 |
| | C | -1.067 | -0.244 | 2.624 | -1.100 | -0.134 | 2.660 | -1.165 | -0.058 | 2.700 | -1.216 | -0.013 | 2.700 | -1.231 | 0.042 | 2.714 |
| | C | -0.656 | 1.134 | 2.056 | -0.702 | 1.251 | 2.094 | -0.696 | 1.288 | 2.110 | -0.726 | 1.310 | 2.073 | -0.722 | 1.319 | 2.005 |
| | C | -1.664 | 2.217 | 2.496 | -1.734 | 2.321 | 2.495 | -1.669 | 2.419 | 2.493 | -1.697 | 2.456 | 2.414 | -1.687 | 2.489 | 2.285 |
| | C | 0.705 | 1.546 | 2.684 | 0.656 | 1.660 | 2.720 | 0.680 | 1.646 | 2.721 | 0.645 | 1.675 | 2.701 | 0.640 | 1.718 | 2.641 |
| | C | 2.001 | 0.754 | 2.381 | 1.940 | 0.840 | 2.415 | 1.932 | 0.789 | 2.393 | 1.910 | 0.823 | 2.405 | 1.924 | 0.885 | 2.395 |
| | C | 2.281 | 0.593 | 0.909 | 2.202 | 0.658 | 0.942 | 2.157 | 0.625 | 0.915 | 2.178 | 0.642 | 0.934 | 2.248 | 0.673 | 0.938 |
| | C | 2.077 | -0.577 | 0.259 | 1.958 | -0.513 | 0.296 | 1.867 | -0.536 | 0.260 | 1.920 | -0.528 | 0.290 | 2.064 | -0.522 | 0.329 |

| | | | | | | | | | | | | | | | | |
|---|---|---|---|---|---|---|---|---|---|---|---|---|---|---|---|---|
| | C | 1.910 | -0.608 | -1.249 | 1.792 | -0.548 | -1.212 | 1.694 | -0.553 | -1.246 | 1.752 | -0.565 | -1.218 | 1.935 | -0.616 | -1.181 |
| | C | 1.440 | 0.726 | -1.899 | 1.402 | 0.802 | -1.881 | 1.356 | 0.815 | -1.905 | 1.403 | 0.791 | -1.897 | 1.521 | 0.695 | -1.909 |
| | C | 1.864 | -1.919 | 0.914 | 1.744 | -1.853 | 0.955 | 1.669 | -1.881 | 0.911 | 1.702 | -1.864 | 0.955 | 1.815 | -1.835 | 1.029 |
| | C | 2.574 | 1.843 | 0.057 | 2.544 | 1.891 | 0.084 | 2.531 | 1.855 | 0.070 | 2.541 | 1.867 | 0.075 | 2.565 | 1.889 | 0.047 |
| | C | 3.925 | 1.656 | -0.679 | 3.896 | 1.648 | -0.634 | 3.876 | 1.580 | -0.652 | 3.893 | 1.605 | -0.636 | 3.941 | 1.677 | -0.634 |
| | C | 2.735 | 3.166 | 0.828 | 2.741 | 3.209 | 0.852 | 2.766 | 3.164 | 0.845 | 2.752 | 3.187 | 0.839 | 2.693 | 3.243 | 0.769 |
| | H | -3.767 | 0.966 | -1.270 | -3.546 | 0.693 | -1.432 | -3.571 | 0.781 | -1.332 | -3.500 | 0.642 | -1.476 | -3.745 | 0.740 | -1.277 |
| | H | 1.567 | 2.843 | -1.565 | 1.586 | 2.917 | -1.553 | 1.591 | 2.922 | -1.553 | 1.599 | 2.904 | -1.567 | 1.594 | 2.823 | -1.638 |
| | H | -0.655 | 2.490 | -1.108 | -0.661 | 2.643 | -1.079 | -0.659 | 2.703 | -1.049 | -0.652 | 2.664 | -1.081 | -0.637 | 2.416 | -1.247 |
| | H | 0.067 | 3.069 | 0.350 | 0.100 | 3.189 | 0.377 | 0.139 | 3.194 | 0.404 | 0.125 | 3.166 | 0.374 | 0.027 | 3.096 | 0.189 |
| | H | -0.056 | 0.115 | 0.280 | 0.024 | 0.289 | 0.369 | 0.004 | 0.282 | 0.394 | -0.001 | 0.257 | 0.389 | -0.050 | 0.135 | 0.353 |
| | H | -2.211 | 1.359 | -2.034 | -1.936 | 0.789 | -2.181 | -2.038 | 0.379 | -2.131 | -1.925 | 0.206 | -2.227 | -1.934 | -0.116 | -2.682 |
| | H | -2.884 | 2.428 | -0.766 | -2.572 | 2.186 | -1.257 | -2.302 | 2.055 | -1.578 | -2.208 | 1.917 | -1.728 | -2.438 | 2.028 | -1.549 |
| | H | -3.959 | -0.304 | 0.642 | -3.607 | -0.717 | 0.313 | -3.499 | -0.784 | 0.253 | -3.508 | -0.814 | 0.240 | -3.587 | -0.855 | 0.340 |
| | H | -2.638 | -1.758 | 2.375 | -2.591 | -1.715 | 2.410 | -2.767 | -1.530 | 2.518 | -2.834 | -1.467 | 2.541 | -2.864 | -1.403 | 2.660 |
| | H | -3.223 | -0.173 | 2.891 | -3.278 | -0.097 | 2.588 | -3.320 | 0.153 | 2.446 | -3.362 | 0.219 | 2.398 | -3.380 | 0.280 | 2.438 |
| | H | -0.341 | -0.996 | 2.281 | -0.310 | -0.859 | 2.417 | -0.405 | -0.828 | 2.502 | -0.460 | -0.795 | 2.540 | -0.486 | -0.759 | 2.589 |
| | H | -0.992 | -0.224 | 3.723 | -1.134 | -0.067 | 3.759 | -1.228 | 0.048 | 3.796 | -1.297 | 0.127 | 3.791 | -1.298 | 0.243 | 3.796 |
| | H | -2.683 | 2.044 | 2.121 | -2.739 | 2.111 | 2.094 | -2.686 | 2.247 | 2.106 | -2.708 | 2.288 | 2.010 | -2.695 | 2.320 | 1.878 |
| | H | -1.721 | 2.274 | 3.595 | -1.821 | 2.393 | 3.591 | -1.736 | 2.513 | 3.589 | -1.787 | 2.573 | 3.507 | -1.786 | 2.653 | 3.370 |
| | H | -1.354 | 3.209 | 2.133 | -1.438 | 3.314 | 2.121 | -1.330 | 3.389 | 2.098 | -1.343 | 3.415 | 2.008 | -1.317 | 3.428 | 1.845 |
| | H | 0.565 | 1.513 | 3.777 | 0.513 | 1.623 | 3.812 | 0.547 | 1.596 | 3.813 | 0.496 | 1.639 | 3.792 | 0.477 | 1.718 | 3.731 |
| | H | 0.867 | 2.605 | 2.457 | 0.830 | 2.718 | 2.493 | 0.888 | 2.700 | 2.509 | 0.853 | 2.727 | 2.482 | 0.834 | 2.767 | 2.391 |
| | H | 1.942 | -0.225 | 2.869 | 1.871 | -0.131 | 2.917 | 1.838 | -0.186 | 2.880 | 1.809 | -0.149 | 2.899 | 1.826 | -0.079 | 2.906 |
| | H | 2.820 | 1.287 | 2.892 | 2.771 | 1.368 | 2.910 | 2.789 | 1.284 | 2.877 | 2.750 | 1.326 | 2.910 | 2.740 | 1.413 | 2.917 |
| | H | 1.169 | -1.398 | -1.468 | 1.012 | -1.299 | -1.422 | 0.890 | -1.276 | -1.459 | 0.952 | -1.297 | -1.419 | 1.181 | -1.399 | -1.378 |
| | H | 2.841 | -0.957 | -1.731 | 2.706 | -0.955 | -1.683 | 2.594 | -0.991 | -1.716 | 2.653 | -1.004 | -1.685 | 2.868 | -1.013 | -1.623 |
| | H | 2.121 | 0.967 | -2.730 | 2.124 | 1.013 | -2.684 | 2.093 | 1.011 | -2.699 | 2.146 | 0.989 | -2.684 | 2.256 | 0.910 | -2.700 |
| | H | 0.452 | 0.583 | -2.359 | 0.433 | 0.688 | -2.386 | 0.388 | 0.741 | -2.418 | 0.441 | 0.702 | -2.419 | 0.560 | 0.550 | -2.425 |
| | H | 2.087 | -1.930 | 1.987 | 1.952 | -1.855 | 2.032 | 1.892 | -1.889 | 1.985 | 1.931 | -1.864 | 2.027 | 2.056 | -1.815 | 2.097 |
| | H | 0.826 | -2.277 | 0.774 | 0.715 | -2.219 | 0.790 | 0.641 | -2.252 | 0.760 | 0.665 | -2.215 | 0.818 | 0.760 | -2.151 | 0.919 |
| | H | 2.517 | -2.675 | 0.442 | 2.417 | -2.599 | 0.496 | 2.346 | -2.613 | 0.437 | 2.361 | -2.617 | 0.487 | 2.438 | -2.628 | 0.576 |
| | H | 4.114 | 2.489 | -1.375 | 4.121 | 2.468 | -1.336 | 4.112 | 2.393 | -1.358 | 4.126 | 2.416 | -1.345 | 4.160 | 2.488 | -1.348 |
| | H | 4.745 | 1.638 | 0.056 | 4.707 | 1.608 | 0.110 | 4.690 | 1.529 | 0.088 | 4.702 | 1.566 | 0.111 | 4.733 | 1.676 | 0.131 |
| | H | 3.990 | 0.722 | -1.250 | 3.931 | 0.707 | -1.195 | 3.888 | 0.636 | -1.210 | 3.919 | 0.658 | -1.187 | 4.020 | 0.726 | -1.173 |
| | H | 3.117 | 3.937 | 0.142 | 3.142 | 3.969 | 0.164 | 3.181 | 3.915 | 0.157 | 3.170 | 3.936 | 0.149 | 3.124 | 3.982 | 0.077 |
| | H | 1.809 | 3.567 | 1.250 | 1.829 | 3.638 | 1.278 | 1.868 | 3.615 | 1.278 | 1.843 | 3.633 | 1.254 | 1.748 | 3.670 | 1.116 |
| | H | 3.460 | 3.060 | 1.649 | 3.464 | 3.081 | 1.669 | 3.491 | 3.017 | 1.658 | 3.466 | 3.054 | 1.663 | 3.366 | 3.164 | 1.635 |
| 4 | C | 2.485 | 4.468 | -3.778 | 2.480 | 4.424 | -3.667 | 2.468 | 4.544 | -3.672 | 2.445 | 4.607 | -3.697 | 2.444 | 4.646 | -4.020 |

| | | | | | | | | | | | | | | | |
|---|---|---|---|---|---|---|---|---|---|---|---|---|---|---|---|
| H | 2.569 | 3.692 | -3.006 | 2.621 | 3.652 | -2.898 | 2.682 | 3.847 | -2.851 | 2.676 | 3.952 | -2.847 | 2.510 | 3.945 | -3.178 |
| H | 3.039 | 5.356 | -3.439 | 3.016 | 5.332 | -3.350 | 2.962 | 5.503 | -3.450 | 2.914 | 5.586 | -3.523 | 2.938 | 5.591 | -3.750 |
| C | 1.042 | 4.864 | -4.001 | 1.001 | 4.759 | -3.832 | 0.956 | 4.756 | -3.768 | 0.926 | 4.762 | -3.770 | 0.986 | 4.865 | -4.329 |
| O | 0.694 | 5.827 | -4.647 | 0.670 | 5.762 | -4.474 | 0.522 | 5.721 | -4.436 | 0.443 | 5.741 | -4.410 | 0.725 | 5.977 | -5.034 |
| O | 0.192 | 4.059 | -3.352 | 0.183 | 3.962 | -3.221 | 0.231 | 3.952 | -3.099 | 0.234 | 3.932 | -3.124 | 0.109 | 4.121 | -3.955 |
| H | -0.763 | 4.375 | -3.371 | -1.071 | 4.265 | -2.715 | -1.099 | 4.129 | -2.248 | -1.119 | 4.085 | -2.227 | -1.111 | 4.060 | -2.722 |
| C | 0.307 | 0.850 | -3.518 | 0.327 | 0.832 | -3.566 | 0.314 | 0.817 | -3.561 | 0.315 | 0.813 | -3.559 | 0.084 | 0.856 | -3.220 |
| H | 1.029 | 1.636 | -3.783 | 1.018 | 1.632 | -3.867 | 0.988 | 1.629 | -3.865 | 0.989 | 1.623 | -3.866 | 0.782 | 1.689 | -3.393 |
| H | -0.664 | 1.167 | -3.936 | -0.660 | 1.130 | -3.960 | -0.689 | 1.099 | -3.926 | -0.687 | 1.097 | -3.925 | -0.879 | 1.198 | -3.633 |
| C | 0.125 | 0.765 | -2.014 | 0.188 | 0.763 | -2.055 | 0.212 | 0.718 | -2.049 | 0.215 | 0.711 | -2.047 | -0.075 | 0.602 | -1.729 |
| O | 0.104 | -0.240 | -1.355 | 0.093 | -0.248 | -1.405 | 0.091 | -0.309 | -1.427 | 0.094 | -0.315 | -1.425 | 0.290 | -0.394 | -1.161 |
| O | -0.105 | 1.962 | -1.429 | 0.086 | 1.963 | -1.467 | 0.177 | 1.902 | -1.428 | 0.184 | 1.894 | -1.421 | -0.656 | 1.591 | -1.024 |
| H | 0.008 | 2.703 | -2.059 | 0.243 | 2.707 | -2.113 | 0.340 | 2.660 | -2.054 | 0.347 | 2.643 | -2.049 | -1.071 | 2.339 | -1.531 |
| C | -0.936 | 8.870 | -1.490 | -0.843 | 8.897 | -1.498 | -0.828 | 8.914 | -1.495 | -0.821 | 8.922 | -1.492 | -0.849 | 8.681 | -1.538 |
| C | -2.299 | 8.360 | -2.036 | -2.215 | 8.417 | -2.048 | -2.204 | 8.454 | -2.046 | -2.191 | 8.451 | -2.047 | -2.208 | 8.136 | -2.057 |
| C | -2.714 | 6.909 | -1.672 | -2.676 | 6.999 | -1.626 | -2.669 | 7.035 | -1.659 | -2.647 | 7.027 | -1.658 | -2.622 | 6.705 | -1.638 |
| C | -2.928 | 5.966 | -2.878 | -2.763 | 5.982 | -2.715 | -2.723 | 6.029 | -2.717 | -2.687 | 6.019 | -2.733 | -2.964 | 5.773 | -2.784 |
| O | -2.181 | 4.715 | -2.695 | -1.835 | 4.483 | -2.000 | -1.853 | 4.199 | -1.599 | -1.870 | 4.156 | -1.590 | -1.722 | 3.841 | -1.993 |
| C | -2.852 | 6.485 | -4.293 | -2.355 | 6.242 | -4.112 | -2.310 | 6.227 | -4.080 | -2.226 | 6.206 | -4.057 | -3.238 | 6.134 | -4.046 |
| C | -3.617 | 4.672 | -2.661 | -3.209 | 4.609 | -2.382 | -3.016 | 4.599 | -2.285 | -3.010 | 4.593 | -2.296 | -3.032 | 4.307 | -2.377 |
| C | -4.269 | 4.317 | -1.351 | -4.170 | 4.404 | -1.236 | -4.201 | 4.488 | -1.338 | -4.204 | 4.496 | -1.360 | -3.986 | 4.107 | -1.200 |
| C | -3.934 | 5.279 | -0.204 | -4.007 | 5.445 | -0.127 | -4.081 | 5.481 | -0.188 | -4.081 | 5.483 | -0.205 | -3.802 | 5.127 | -0.071 |
| C | -3.882 | 6.764 | -0.636 | -3.899 | 6.901 | -0.638 | -3.908 | 6.939 | -0.660 | -3.894 | 6.940 | -0.676 | -3.777 | 6.595 | -0.560 |
| C | -5.219 | 7.166 | -1.286 | -5.185 | 7.302 | -1.383 | -5.161 | 7.405 | -1.425 | -5.147 | 7.411 | -1.438 | -5.134 | 6.957 | -1.198 |
| C | -3.739 | 7.659 | 0.627 | -3.753 | 7.834 | 0.590 | -3.751 | 7.847 | 0.582 | -3.742 | 7.846 | 0.572 | -3.628 | 7.546 | 0.657 |
| C | -2.465 | 7.638 | 1.510 | -2.486 | 7.792 | 1.490 | -2.487 | 7.779 | 1.484 | -2.482 | 7.782 | 1.480 | -2.344 | 7.581 | 1.525 |
| C | -1.185 | 7.838 | 0.738 | -1.188 | 7.918 | 0.736 | -1.199 | 7.905 | 0.719 | -1.186 | 7.915 | 0.726 | -1.077 | 7.743 | 0.727 |
| C | -0.367 | 6.802 | 0.439 | -0.402 | 6.842 | 0.473 | -0.419 | 6.822 | 0.447 | -0.404 | 6.836 | 0.457 | -0.266 | 6.692 | 0.462 |
| C | 0.622 | 6.906 | -0.705 | 0.607 | 6.880 | -0.659 | 0.590 | 6.866 | -0.682 | 0.604 | 6.877 | -0.675 | 0.709 | 6.748 | -0.696 |
| C | 0.249 | 7.932 | -1.816 | 0.322 | 7.923 | -1.778 | 0.326 | 7.931 | -1.784 | 0.341 | 7.945 | -1.775 | 0.329 | 7.729 | -1.842 |
| C | -0.414 | 5.432 | 1.067 | -0.489 | 5.500 | 1.157 | -0.487 | 5.492 | 1.155 | -0.486 | 5.499 | 1.149 | -0.303 | 5.347 | 1.148 |
| C | -0.927 | 9.184 | 0.034 | -0.861 | 9.242 | 0.018 | -0.858 | 9.237 | 0.024 | -0.849 | 9.246 | 0.028 | -0.826 | 9.062 | -0.033 |
| C | 0.450 | 9.753 | 0.460 | 0.524 | 9.762 | 0.477 | 0.529 | 9.735 | 0.504 | 0.537 | 9.751 | 0.502 | 0.555 | 9.649 | 0.361 |
| C | -1.919 | 10.313 | 0.374 | -1.821 | 10.413 | 0.303 | -1.808 | 10.413 | 0.320 | -1.804 | 10.418 | 0.324 | -1.821 | 10.204 | 0.255 |
| H | -3.579 | 7.294 | -4.457 | -2.749 | 7.218 | -4.434 | -2.381 | 7.255 | -4.454 | -2.226 | 7.230 | -4.450 | -3.256 | 7.176 | -4.375 |
| H | -0.757 | 9.822 | -2.022 | -0.631 | 9.833 | -2.046 | -0.605 | 9.854 | -2.031 | -0.602 | 9.863 | -2.027 | -0.678 | 9.610 | -2.112 |
| H | -2.242 | 8.469 | -3.126 | -2.150 | 8.487 | -3.141 | -2.152 | 8.546 | -3.138 | -2.144 | 8.548 | -3.139 | -2.133 | 8.180 | -3.152 |
| H | -3.087 | 9.070 | -1.746 | -2.986 | 9.151 | -1.779 | -2.967 | 9.186 | -1.753 | -2.961 | 9.176 | -1.752 | -3.004 | 8.854 | -1.812 |
| H | -1.855 | 6.447 | -1.188 | -1.852 | 6.597 | -1.023 | -1.861 | 6.596 | -1.023 | -1.844 | 6.589 | -1.029 | -1.760 | 6.231 | -1.165 |

| | | | | | | | | | | | | | | | | |
|---|---|---|---|---|---|---|---|---|---|---|---|---|---|---|---|---|
| | H | -3.076 | 5.672 | -4.999 | -2.723 | 5.458 | -4.786 | -2.775 | 5.497 | -4.757 | -2.609 | 5.462 | -4.767 | -3.515 | 5.387 | -4.791 |
| | H | -1.850 | 6.869 | -4.527 | -1.250 | 6.279 | -4.229 | -1.177 | 5.966 | -4.153 | -0.997 | 5.929 | -4.126 | -0.244 | 6.039 | -5.150 |
| | H | -4.057 | 4.171 | -3.532 | -3.473 | 3.986 | -3.248 | -3.195 | 3.969 | -3.173 | -3.194 | 3.966 | -3.186 | -3.390 | 3.700 | -3.223 |
| | H | -3.990 | 3.278 | -1.096 | -4.013 | 3.381 | -0.854 | -4.192 | 3.450 | -0.968 | -4.209 | 3.455 | -0.995 | -3.847 | 3.075 | -0.832 |
| | H | -5.357 | 4.319 | -1.543 | -5.188 | 4.437 | -1.658 | -5.138 | 4.630 | -1.898 | -5.135 | 4.654 | -1.926 | -5.010 | 4.173 | -1.605 |
| | H | -2.968 | 5.005 | 0.245 | -3.114 | 5.209 | 0.468 | -3.230 | 5.198 | 0.449 | -3.232 | 5.192 | 0.431 | -2.869 | 4.917 | 0.471 |
| | H | -4.690 | 5.158 | 0.590 | -4.866 | 5.379 | 0.560 | -4.981 | 5.426 | 0.446 | -4.982 | 5.435 | 0.428 | -4.624 | 5.009 | 0.656 |
| | H | -6.059 | 6.956 | -0.604 | -6.072 | 7.180 | -0.741 | -6.060 | 7.307 | -0.797 | -6.048 | 7.296 | -0.815 | -5.948 | 6.852 | -0.461 |
| | H | -5.245 | 8.243 | -1.514 | -5.150 | 8.352 | -1.710 | -5.084 | 8.460 | -1.730 | -5.075 | 8.471 | -1.726 | -5.143 | 8.000 | -1.551 |
| | H | -5.411 | 6.630 | -2.227 | -5.343 | 6.691 | -2.286 | -5.324 | 6.810 | -2.337 | -5.303 | 6.837 | -2.365 | -5.383 | 6.328 | -2.062 |
| | H | -4.580 | 7.390 | 1.286 | -4.597 | 7.591 | 1.250 | -4.597 | 7.594 | 1.237 | -4.591 | 7.591 | 1.223 | -4.455 | 7.295 | 1.340 |
| | H | -3.946 | 8.690 | 0.322 | -3.936 | 8.861 | 0.256 | -3.926 | 8.883 | 0.272 | -3.919 | 8.882 | 0.264 | -3.853 | 8.559 | 0.305 |
| | H | -2.429 | 6.689 | 2.053 | -2.495 | 6.865 | 2.072 | -2.500 | 6.841 | 2.046 | -2.495 | 6.843 | 2.041 | -2.295 | 6.668 | 2.126 |
| | H | -2.607 | 8.410 | 2.286 | -2.608 | 8.602 | 2.228 | -2.597 | 8.578 | 2.234 | -2.602 | 8.578 | 2.232 | -2.478 | 8.402 | 2.248 |
| | H | 0.699 | 5.901 | -1.155 | 0.626 | 5.869 | -1.101 | 0.599 | 5.863 | -1.137 | 0.603 | 5.872 | -1.129 | 0.776 | 5.726 | -1.106 |
| | H | 1.634 | 7.117 | -0.314 | 1.623 | 7.028 | -0.252 | 1.606 | 6.988 | -0.268 | 1.623 | 6.997 | -0.266 | 1.727 | 6.969 | -0.327 |
| | H | 1.133 | 8.555 | -2.010 | 1.231 | 8.522 | -1.922 | 1.242 | 8.524 | -1.909 | 1.255 | 8.543 | -1.893 | 1.212 | 8.337 | -2.084 |
| | H | 0.052 | 7.401 | -2.759 | 0.167 | 7.403 | -2.734 | 0.170 | 7.429 | -2.750 | 0.191 | 7.449 | -2.743 | 0.103 | 7.163 | -2.759 |
| | H | -0.744 | 4.670 | 0.337 | -0.856 | 4.721 | 0.465 | -0.879 | 4.702 | 0.491 | -0.882 | 4.723 | 0.473 | -0.685 | 4.564 | 0.467 |
| | H | 0.597 | 5.131 | 1.389 | 0.520 | 5.179 | 1.468 | 0.531 | 5.182 | 1.448 | 0.528 | 5.175 | 1.441 | 0.720 | 5.035 | 1.421 |
| | H | -1.062 | 5.365 | 1.949 | -1.116 | 5.498 | 2.057 | -1.090 | 5.513 | 2.071 | -1.091 | 5.513 | 2.064 | -0.900 | 5.334 | 2.068 |
| | H | 1.281 | 9.047 | 0.340 | 1.325 | 9.017 | 0.398 | 1.323 | 8.984 | 0.421 | 1.332 | 9.000 | 0.419 | 1.381 | 8.931 | 0.283 |
| | H | 0.687 | 10.655 | -0.129 | 0.817 | 10.643 | -0.120 | 0.836 | 10.619 | -0.079 | 0.840 | 10.635 | -0.083 | 0.796 | 10.522 | -0.272 |
| | H | 0.421 | 10.048 | 1.521 | 0.474 | 10.078 | 1.531 | 0.472 | 10.040 | 1.560 | 0.483 | 10.055 | 1.560 | 0.526 | 9.998 | 1.406 |
| | H | -1.963 | 10.481 | 1.461 | -1.900 | 10.609 | 1.383 | -1.897 | 10.588 | 1.402 | -1.889 | 10.598 | 1.406 | -1.872 | 10.426 | 1.331 |
| | H | -1.574 | 11.247 | -0.096 | -1.424 | 11.324 | -0.171 | -1.394 | 11.328 | -0.131 | -1.396 | 11.334 | -0.133 | -1.473 | 11.119 | -0.252 |
| | H | -2.942 | 10.142 | 0.023 | -2.836 | 10.278 | -0.085 | -2.820 | 10.297 | -0.081 | -2.816 | 10.297 | -0.074 | -2.841 | 10.017 | -0.097 |
| 5 | C | 1.224 | 5.668 | -1.673 | 1.231 | 5.632 | -1.611 | 1.321 | 5.719 | -1.596 | 1.331 | 5.784 | -1.553 | 1.322 | 5.793 | -1.629 |
| | H | 0.969 | 4.614 | -1.501 | 0.970 | 4.578 | -1.438 | 1.207 | 4.642 | -1.411 | 1.275 | 4.708 | -1.339 | 1.254 | 4.718 | -1.424 |
| | H | 2.268 | 5.829 | -1.363 | 2.281 | 5.793 | -1.321 | 2.346 | 6.022 | -1.330 | 2.335 | 6.149 | -1.293 | 2.318 | 6.155 | -1.335 |
| | C | 0.334 | 6.573 | -0.842 | 0.345 | 6.547 | -0.757 | 0.311 | 6.480 | -0.725 | 0.275 | 6.479 | -0.682 | 0.244 | 6.463 | -0.808 |
| | O | 0.511 | 7.760 | -0.705 | 0.686 | 7.716 | -0.569 | 0.522 | 7.684 | -0.477 | 0.457 | 7.686 | -0.358 | 0.426 | 7.761 | -0.602 |
| | O | -0.637 | 5.887 | -0.220 | -0.697 | 5.976 | -0.238 | -0.657 | 5.794 | -0.255 | -0.686 | 5.774 | -0.272 | -0.694 | 5.850 | -0.327 |
| | H | -1.135 | 6.404 | 0.486 | -1.283 | 6.222 | 1.035 | -1.358 | 5.753 | 1.216 | -1.375 | 5.665 | 1.252 | -1.498 | 5.719 | 1.369 |
| | C | -2.664 | 4.115 | -2.044 | -2.631 | 4.211 | -2.089 | -2.674 | 4.151 | -2.093 | -2.688 | 4.137 | -2.090 | -2.723 | 4.093 | -2.077 |
| | H | -1.718 | 4.604 | -2.317 | -1.703 | 4.720 | -2.384 | -1.753 | 4.690 | -2.354 | -1.768 | 4.682 | -2.342 | -1.786 | 4.615 | -2.317 |
| | H | -3.270 | 4.879 | -1.527 | -3.244 | 4.980 | -1.586 | -3.308 | 4.874 | -1.549 | -3.327 | 4.853 | -1.545 | -3.353 | 4.821 | -1.537 |
| | C | -2.428 | 2.978 | -1.065 | -2.354 | 3.122 | -1.065 | -2.378 | 3.005 | -1.137 | -2.400 | 2.983 | -1.144 | -2.470 | 2.921 | -1.145 |
| | O | -2.888 | 1.870 | -1.148 | -2.853 | 2.022 | -1.067 | -2.878 | 1.908 | -1.192 | -2.918 | 1.894 | -1.195 | -2.921 | 1.812 | -1.261 |

| | | | | | | | | | | | | | | | |
|---|---|---|---|---|---|---|---|---|---|---|---|---|---|---|---|
| O | -1.674 | 3.315 | 0.002 | -1.536 | 3.498 | -0.075 | -1.543 | 3.328 | -0.143 | -1.546 | 3.286 | -0.159 | -1.705 | 3.235 | -0.080 |
| H | -1.299 | 4.219 | -0.072 | -1.168 | 4.425 | -0.201 | -1.144 | 4.240 | -0.255 | -1.152 | 4.193 | -0.271 | -1.354 | 4.145 | -0.142 |
| C | 2.273 | 7.731 | 3.617 | 2.328 | 7.908 | 3.678 | 2.329 | 7.785 | 3.646 | 2.347 | 7.781 | 3.638 | 2.320 | 7.722 | 3.615 |
| C | 0.866 | 8.323 | 3.908 | 0.935 | 8.535 | 3.959 | 0.954 | 8.432 | 3.964 | 0.965 | 8.421 | 3.933 | 0.913 | 8.321 | 3.890 |
| C | -0.339 | 7.344 | 3.875 | -0.276 | 7.569 | 3.941 | -0.279 | 7.500 | 3.926 | -0.261 | 7.483 | 3.886 | -0.306 | 7.368 | 3.899 |
| C | -1.439 | 7.722 | 2.860 | -1.267 | 7.796 | 2.850 | -1.239 | 7.712 | 2.849 | -1.220 | 7.695 | 2.785 | -1.420 | 7.723 | 2.929 |
| O | -1.754 | 6.591 | 1.981 | -1.499 | 6.081 | 2.069 | -1.664 | 5.548 | 2.143 | -1.688 | 5.488 | 2.171 | -1.870 | 5.452 | 2.222 |
| C | -1.399 | 9.042 | 2.131 | -1.105 | 8.821 | 1.793 | -1.110 | 8.693 | 1.793 | -1.066 | 8.614 | 1.716 | -1.542 | 8.844 | 2.196 |
| C | -2.742 | 7.026 | 2.936 | -2.449 | 6.911 | 2.760 | -2.334 | 6.666 | 2.686 | -2.327 | 6.651 | 2.661 | -2.464 | 6.620 | 2.790 |
| C | -3.096 | 6.096 | 4.065 | -2.999 | 6.284 | 4.018 | -2.994 | 6.273 | 3.998 | -3.003 | 6.316 | 3.982 | -3.050 | 6.227 | 4.141 |
| C | -1.928 | 5.765 | 5.004 | -1.931 | 6.033 | 5.083 | -1.971 | 5.973 | 5.086 | -1.984 | 6.020 | 5.076 | -1.962 | 5.931 | 5.173 |
| C | -0.976 | 6.956 | 5.258 | -0.977 | 7.229 | 5.313 | -0.967 | 7.122 | 5.319 | -0.960 | 7.156 | 5.280 | -0.926 | 7.069 | 5.327 |
| C | -1.756 | 8.152 | 5.840 | -1.755 | 8.460 | 5.811 | -1.681 | 8.374 | 5.860 | -1.659 | 8.429 | 5.794 | -1.610 | 8.338 | 5.873 |
| C | 0.045 | 6.556 | 6.358 | 0.032 | 6.842 | 6.426 | 0.049 | 6.678 | 6.398 | 0.047 | 6.719 | 6.376 | 0.107 | 6.638 | 6.407 |
| C | 1.104 | 5.447 | 6.122 | 1.052 | 5.690 | 6.202 | 1.038 | 5.514 | 6.120 | 1.034 | 5.545 | 6.128 | 1.117 | 5.488 | 6.152 |
| C | 1.918 | 5.640 | 4.870 | 1.868 | 5.843 | 4.943 | 1.829 | 5.703 | 4.855 | 1.848 | 5.708 | 4.872 | 1.920 | 5.652 | 4.886 |
| C | 1.694 | 4.908 | 3.754 | 1.618 | 5.119 | 3.821 | 1.544 | 5.014 | 3.715 | 1.582 | 5.004 | 3.740 | 1.671 | 4.926 | 3.771 |
| C | 2.152 | 5.404 | 2.398 | 2.093 | 5.596 | 2.463 | 2.012 | 5.519 | 2.366 | 2.047 | 5.501 | 2.385 | 2.131 | 5.410 | 2.408 |
| C | 2.339 | 6.944 | 2.287 | 2.402 | 7.115 | 2.355 | 2.382 | 7.026 | 2.303 | 2.411 | 7.009 | 2.305 | 2.393 | 6.936 | 2.291 |
| C | 0.902 | 3.629 | 3.679 | 0.794 | 3.859 | 3.749 | 0.746 | 3.739 | 3.633 | 0.758 | 3.745 | 3.662 | 0.806 | 3.696 | 3.692 |
| C | 2.857 | 6.856 | 4.760 | 2.861 | 7.016 | 4.832 | 2.850 | 6.852 | 4.772 | 2.864 | 6.861 | 4.777 | 2.890 | 6.841 | 4.760 |
| C | 4.289 | 6.375 | 4.414 | 4.270 | 6.459 | 4.503 | 4.241 | 6.266 | 4.416 | 4.258 | 6.277 | 4.433 | 4.305 | 6.308 | 4.414 |
| C | 3.027 | 7.687 | 6.047 | 3.060 | 7.852 | 6.111 | 3.082 | 7.641 | 6.073 | 3.088 | 7.671 | 6.068 | 3.091 | 7.688 | 6.031 |
| H | -0.513 | 9.122 | 1.485 | -0.246 | 8.618 | 1.126 | -0.509 | 8.230 | 0.928 | -0.401 | 8.097 | 0.801 | -0.252 | 8.087 | 0.033 |
| H | 2.933 | 8.610 | 3.506 | 3.011 | 8.770 | 3.583 | 3.030 | 8.635 | 3.564 | 3.046 | 8.633 | 3.557 | 2.987 | 8.600 | 3.519 |
| H | 0.720 | 9.105 | 3.151 | 0.797 | 9.322 | 3.209 | 0.825 | 9.249 | 3.245 | 0.835 | 9.238 | 3.212 | 0.765 | 9.088 | 3.116 |
| H | 0.898 | 8.875 | 4.859 | 0.965 | 9.074 | 4.916 | 1.011 | 8.940 | 4.936 | 1.005 | 8.929 | 4.906 | 0.948 | 8.894 | 4.827 |
| H | 0.029 | 6.401 | 3.469 | 0.136 | 6.596 | 3.645 | 0.096 | 6.488 | 3.622 | 0.108 | 6.475 | 3.597 | 0.016 | 6.390 | 3.535 |
| H | -1.382 | 9.878 | 2.846 | -0.918 | 9.795 | 2.274 | -0.549 | 9.599 | 2.059 | -0.471 | 9.515 | 1.913 | -0.859 | 9.695 | 2.275 |
| H | -2.290 | 9.143 | 1.496 | -2.002 | 8.889 | 1.163 | -2.084 | 8.920 | 1.334 | -2.000 | 8.808 | 1.170 | -2.414 | 8.972 | 1.547 |
| H | -3.605 | 7.517 | 2.474 | -3.249 | 7.277 | 2.106 | -3.104 | 7.046 | 1.998 | -3.089 | 7.011 | 1.952 | -3.285 | 6.971 | 2.143 |
| H | -3.555 | 5.183 | 3.646 | -3.517 | 5.354 | 3.732 | -3.627 | 5.397 | 3.786 | -3.653 | 5.448 | 3.789 | -3.687 | 5.343 | 3.975 |
| H | -3.898 | 6.597 | 4.633 | -3.780 | 6.961 | 4.399 | -3.671 | 7.080 | 4.312 | -3.666 | 7.144 | 4.271 | -3.710 | 7.037 | 4.486 |
| H | -1.344 | 4.924 | 4.602 | -1.336 | 5.152 | 4.803 | -1.421 | 5.057 | 4.824 | -1.450 | 5.090 | 4.830 | -1.436 | 5.008 | 4.883 |
| H | -2.342 | 5.421 | 5.966 | -2.423 | 5.786 | 6.038 | -2.494 | 5.766 | 6.034 | -2.503 | 5.841 | 6.033 | -2.426 | 5.738 | 6.156 |
| H | -2.237 | 7.872 | 6.793 | -2.295 | 8.233 | 6.746 | -2.160 | 8.151 | 6.826 | -2.178 | 8.227 | 6.745 | -2.059 | 8.141 | 6.859 |
| H | -1.082 | 8.999 | 6.047 | -1.078 | 9.305 | 6.015 | -0.973 | 9.200 | 6.026 | -0.935 | 9.239 | 5.976 | -0.889 | 9.161 | 5.999 |
| H | -2.542 | 8.525 | 5.168 | -2.495 | 8.816 | 5.077 | -2.458 | 8.747 | 5.176 | -2.401 | 8.817 | 5.080 | -2.404 | 8.708 | 5.210 |
| H | -0.551 | 6.232 | 7.226 | -0.576 | 6.558 | 7.302 | -0.556 | 6.370 | 7.267 | -0.566 | 6.433 | 7.248 | -0.485 | 6.339 | 7.288 |

| | | | | | | | | | | | | | | | |
|---|---|---|---|---|---|---|---|---|---|---|---|---|---|---|---|
| H | 0.546 | 7.471 | 6.692 | 0.562 | 7.751 | 6.734 | 0.601 | 7.562 | 6.735 | 0.602 | 7.605 | 6.705 | 0.650 | 7.531 | 6.736 |
| H | 0.599 | 4.478 | 6.091 | 0.515 | 4.736 | 6.193 | 0.477 | 4.576 | 6.071 | 0.471 | 4.607 | 6.094 | 0.574 | 4.539 | 6.137 |
| H | 1.739 | 5.422 | 7.025 | 1.693 | 5.658 | 7.097 | 1.691 | 5.429 | 7.006 | 1.674 | 5.473 | 7.023 | 1.772 | 5.436 | 7.040 |
| H | 1.390 | 5.081 | 1.668 | 1.299 | 5.333 | 1.744 | 1.194 | 5.307 | 1.658 | 1.225 | 5.282 | 1.684 | 1.337 | 5.129 | 1.694 |
| H | 3.078 | 4.884 | 2.090 | 2.969 | 5.004 | 2.138 | 2.858 | 4.901 | 2.008 | 2.892 | 4.885 | 2.025 | 3.023 | 4.847 | 2.076 |
| H | 3.320 | 7.127 | 1.832 | 3.420 | 7.219 | 1.962 | 3.410 | 7.098 | 1.932 | 3.439 | 7.083 | 1.932 | 3.394 | 7.083 | 1.863 |
| H | 1.606 | 7.366 | 1.583 | 1.754 | 7.574 | 1.596 | 1.763 | 7.533 | 1.549 | 1.790 | 7.506 | 1.549 | 1.693 | 7.386 | 1.571 |
| H | -0.031 | 3.767 | 3.100 | -0.114 | 4.007 | 3.138 | -0.168 | 3.878 | 3.031 | -0.173 | 3.912 | 3.094 | -0.136 | 3.913 | 3.155 |
| H | 1.486 | 2.862 | 3.140 | 1.380 | 3.068 | 3.248 | 1.355 | 2.974 | 3.120 | 1.330 | 2.971 | 3.121 | 1.326 | 2.902 | 3.126 |
| H | 0.644 | 3.208 | 4.657 | 0.502 | 3.469 | 4.729 | 0.474 | 3.326 | 4.611 | 0.505 | 3.328 | 4.642 | 0.557 | 3.279 | 4.673 |
| H | 4.949 | 7.241 | 4.236 | 4.968 | 7.287 | 4.292 | 4.954 | 7.084 | 4.217 | 4.969 | 7.091 | 4.210 | 4.991 | 7.145 | 4.203 |
| H | 4.707 | 5.804 | 5.259 | 4.665 | 5.900 | 5.366 | 4.630 | 5.682 | 5.264 | 4.653 | 5.715 | 5.294 | 4.715 | 5.751 | 5.272 |
| H | 4.334 | 5.725 | 3.532 | 4.282 | 5.774 | 3.647 | 4.228 | 5.604 | 3.541 | 4.248 | 5.589 | 3.579 | 4.322 | 5.628 | 3.553 |
| H | 3.296 | 7.046 | 6.898 | 3.281 | 7.210 | 6.976 | 3.313 | 6.966 | 6.909 | 3.300 | 7.010 | 6.920 | 3.332 | 7.056 | 6.898 |
| H | 3.845 | 8.410 | 5.906 | 3.919 | 8.526 | 5.968 | 3.949 | 8.307 | 5.936 | 3.962 | 8.327 | 5.929 | 3.937 | 8.375 | 5.874 |
| H | 2.147 | 8.271 | 6.332 | 2.210 | 8.489 | 6.373 | 2.249 | 8.283 | 6.380 | 2.256 | 8.326 | 6.347 | 2.231 | 8.312 | 6.294 |

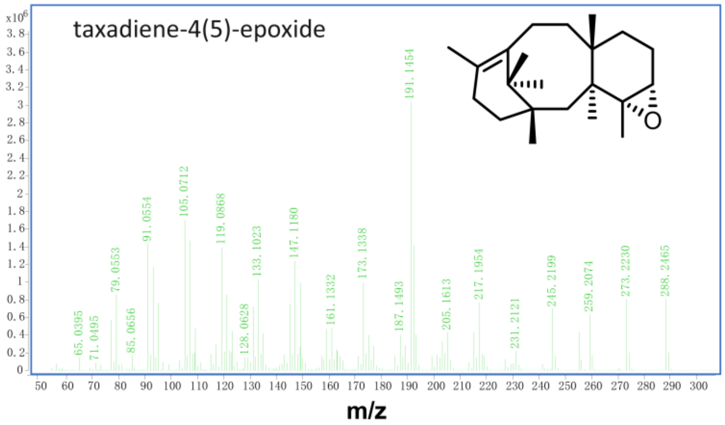

**Figure S1 Mass fragmentation pattern of taxadiene-4(5)-epoxide from GC-MS analysis.**

| | |
|---|---|
| 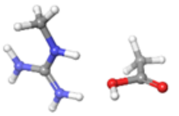 | -0.8005302503E+00 a.u.<br>-0.2178354E+02 eV<br>-0.5023407E+03 kcal/mol |
| 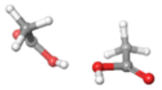 | -0.9134744720E+00 a.u.<br>-0.2485690E+02 eV<br>-0.5732144E+03 kcal/mol |
| 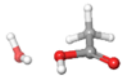 | -0.9328223582E+00 a.u.<br>-0.2538339E+02 eV<br>-0.5853554E+03 kcal/mol |
| 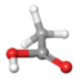 | -0.9479510203E+00 a.u.<br>-0.2579506E+02 eV<br>-0.5948487E+03 kcal/mol |

**Figure S2 Electronic Electrostatic Potential (ESP) analysis.**

Calculated at the B3LYP-D4/def2-SVP level.

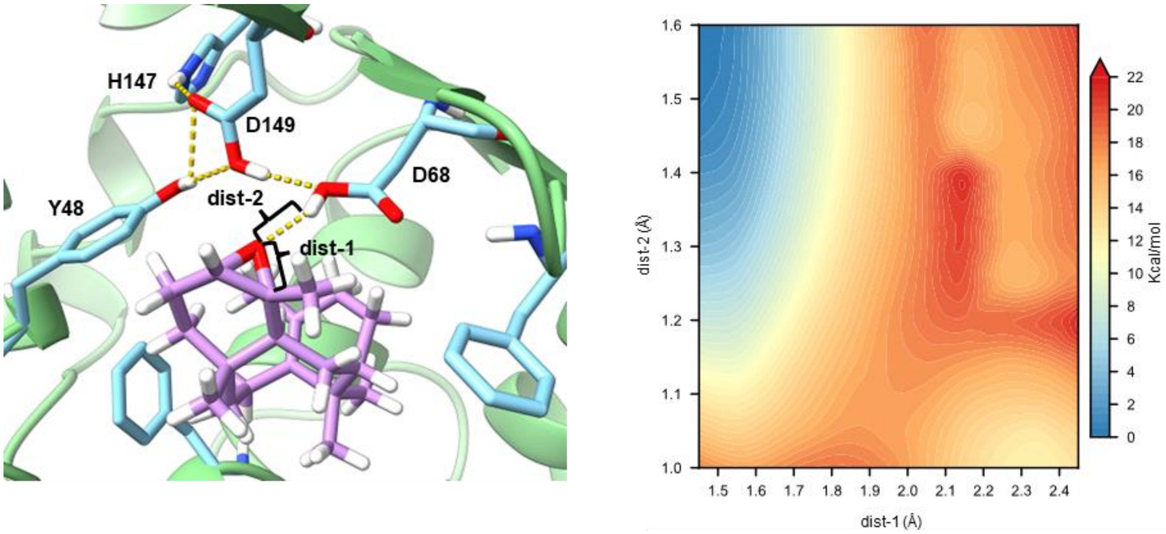

**Figure S3 Two-dimensional potential energy surface (PES) scan for locating the first transition state (TS1).**

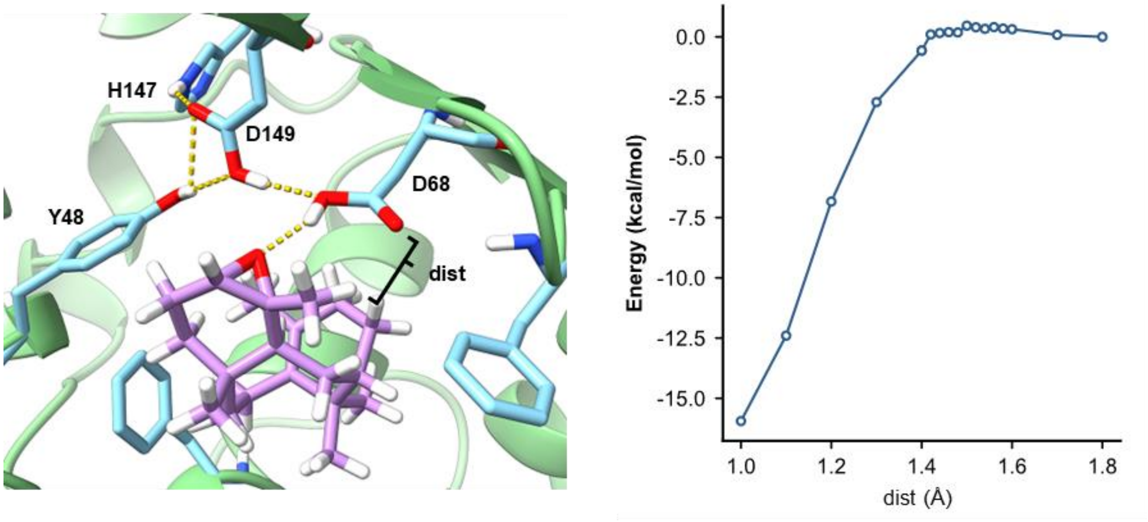

**Figure S4 One-dimensional relaxed potential energy surface (PES) scan for locating the second transition state (TS2).**

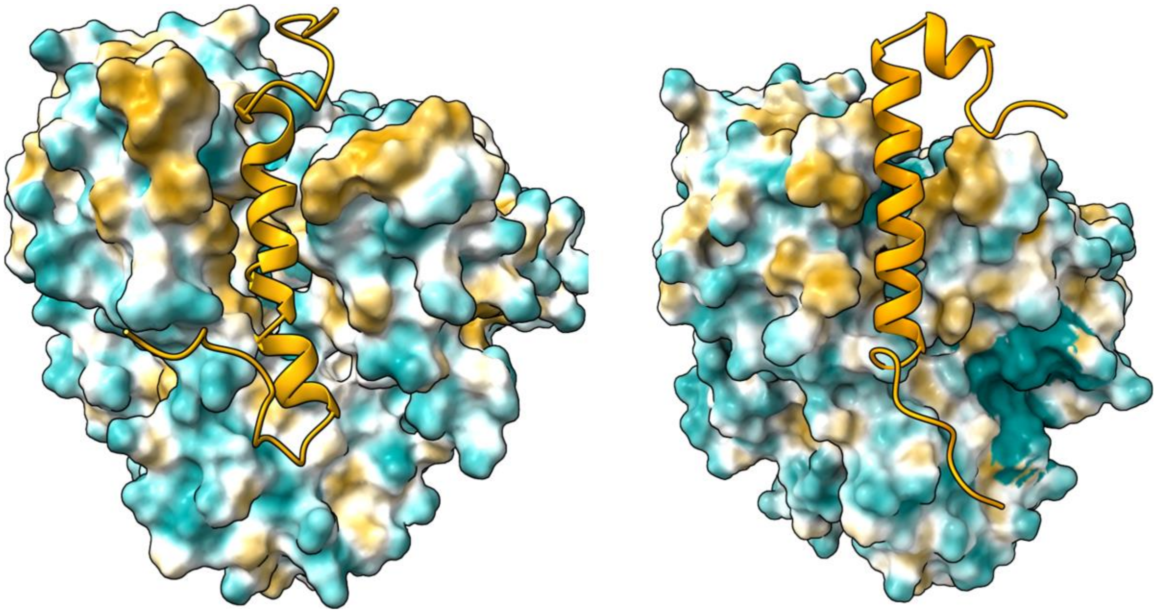

**Figure S5 Structural modeling of the FoTO1 C-terminus interacting with T5alphaOH.** The FoTO1 C-terminal extension is shown binding within the polar surface groove of the T5αOH N-terminus. Binding modes Comp-cn (left) and Comp-cc (right) are displayed to illustrate the spatial complementarity between the two domains.

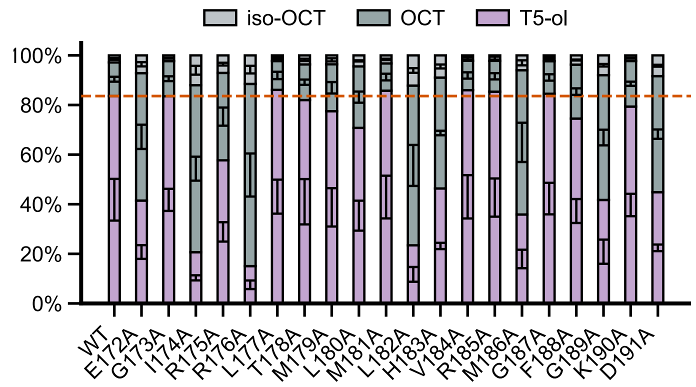

**Figure S6 Alanine scanning mutagenesis across residues 172–191.** Samples that show significant differences from the wild type are marked with an asterisk at the top of the column.
